# Macrophage epigenetic memories of early life injury drive neonatal nociceptive priming

**DOI:** 10.1101/2023.02.13.528015

**Authors:** Adam J. Dourson, Adewale O. Fadaka, Anna M. Warshak, Aditi Paranjpe, Benjamin Weinhaus, Luis F. Queme, Megan C. Hofmann, Heather M. Evans, Omer A. Donmez, Carmy Forney, Matthew T. Weirauch, Leah C. Kottyan, Daniel Lucas, George S. Deepe, Michael P. Jankowski

**Author notes:** Corresponding author: Michael P. Jankowski 3333 Burnet Avenue, MLC6016 Cincinnati, Ohio 45229-3026. Conflict of interest: The authors declare no competing financial interests.

## Abstract

The developing peripheral nervous and immune systems are functionally distinct from adults. These systems are vulnerable to early life injury, which influences outcomes related to nociception following subsequent injury later in life (i.e., “neonatal nociceptive priming”). The underpinnings of this phenomenon are largely unknown, although previous work indicates that macrophages are epigenetically trained by inflammation and injury. We found that macrophages are both necessary and partially sufficient to drive neonatal nociceptive priming possibly due to a long-lasting epigenetic remodeling. The p75 neurotrophic factor receptor (NTR) was an important effector in regulating neonatal nociceptive priming through modulation of the inflammatory profile of rodent and human macrophages. This “pain memory” was long lasting in females and could be transferred to a naïve host to alter sex-specific pain-related behaviors. This study reveals a novel mechanism by which acute, neonatal post-surgical pain drives a peripheral immune-related predisposition to persistent pain following a subsequent injury.

**Graphical Abstract:** 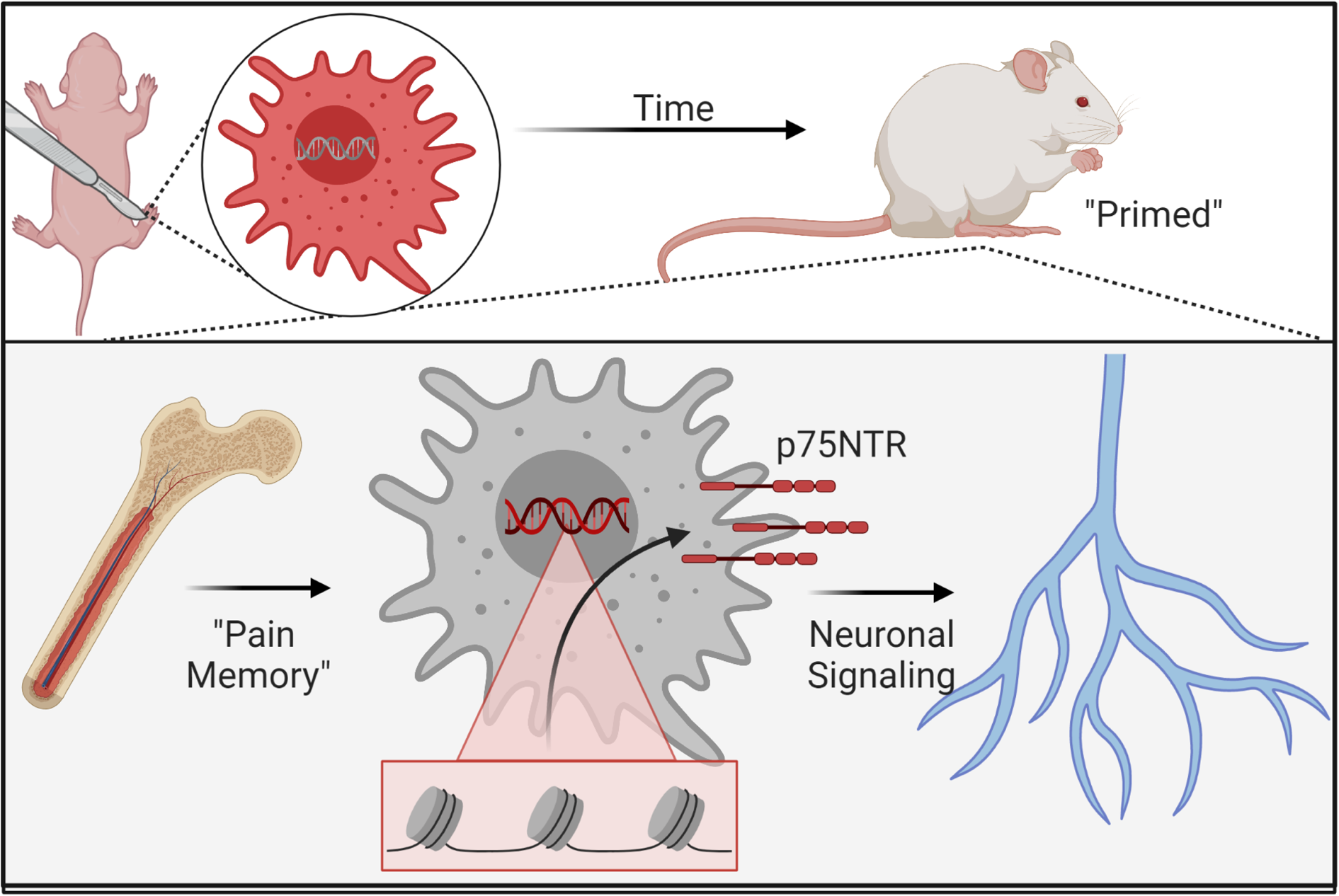

## Introduction

Peripheral injury activates several biological systems to evoke an appropriate motor and affective response to sensory stimuli. To generate and maintain nociceptive signals, there is remarkable crosstalk between cellular systems to create distinct molecular patterns that depend on the given aversive event^1,2^. Importantly, these systems undergo significant developmental changes especially in early life^3,4^, a period in which nociceptors and immune cells display unique properties compared to their adult counterparts^5–10^.

Critical periods are constrained time points of high plasticity in which typical development is persistently altered by external input. Clinical studies indicate that early life surgery or time spent in the neonatal intensive care unit (NICU) alters the development of the somatosensory system^11,12^. In these patients, injury in adulthood has a higher risk of complications including increased pain management medication use and longer hospital stays. Rodent drug studies revealed that the peripheral afferent input is necessary for this phenomenon termed “neonatal nociceptive priming”^13^. However, other peripheral and developmental factors that influence this experience are unknown.

Immune cells, particularly macrophages, modulate nociceptive responses. For example, patient recovery from surgery correlates with their pre-operative inflammatory profile^14–20^. In rodents, the depletion or activation/transfer of macrophages results in differential pain-like behaviors. These studies indicate a key role for neuroimmune signaling in the nociceptive response^5,20–23^. Evidence also suggests that macrophages “learn” in response to multiple exposures of the same or similar adverse agents, likely through epigenetic remodeling^24–26^. While these studies indicate a cellular “memory” of these innate cells, it is unknown if there is a similar effect after injury, or whether this “memory” may persistently affect behavior.

Aversive stimuli within the first week of life alters sensory neuron development by modifying key signaling pathways, such as neurotrophic factors^6,7,27^. Similar effects may be present in immune cells after an injury, however, much less is understood^28,29^. p75NTR (also known as nerve growth factor receptor (NGFR)) modulates broad neurotrophic factor signaling pathways^30^, transcription, and the function of neurons^31^ and immune cells^28^. Neurotrophins, therefore, have critical effects on the development of the nervous and immune systems and could play a dual role in both neonatal inflammation and pain.

Pediatric post-surgical pain is a major clinical health problem that is complicated by underlying differences in patients, diagnostic uncertainty, and developmental concerns^32–35^. There is often a choice between patients suffering the negative effects of pain (intensity or unpleasantness and developmental consequences) or the side effects of current treatments (NSAIDs, steroids, anticonvulsants, opioids, etc.^36,37^). These and other studies clearly demonstrate that improved therapeutics designed for early life pain are necessary for precision care^38^. Here, we aimed to investigate how early life injury alters the peripheral neuroimmune system to “prime” animals to re-injury later in life. We determined the necessity and sufficiency of macrophages for neonatal nociceptive priming, and we evaluated the epigenetic changes that occur in macrophages after neonatal injury, which may underlie this phenomenon. We hypothesized that injury-induced epigenetic changes to chromatin accessibility in developing macrophages contributes to neonatal nociceptive priming.

## Results

### Macrophages are necessary for neonatal nociceptive priming

Previous data indicate that macrophages are important for acute nociception in adults and neonates^5,21,39^. To establish the role of neonatal macrophages in the induction of neonatal nociceptive priming, we used macrophage fas-induced apoptosis (MaFIA) animals to ablate macrophages early in life and tested animals before and after both neonatal and adolescent incision (Fig. 1A). In this animal, *Csf1r*-driven expression of the FK506 binding protein 1A induces cell apoptosis when dimerized, which is temporally controlled by the injection of a designer chemical and inducer of dimerization, AP20187 (AP). Importantly, when injected, *Csf1r*+ cells undergo apoptosis, but within days of stopping the injection, new monocytes derived from the bone marrow begin to replenish the lost cells^40^. We found that systemic injection of AP drastically reduced splenic macrophages but not microglia in the spinal cord (Supp. Fig. 1A-B), as has been similarly noted in adults^41,42^. This knockout strategy also prevented macrophage infiltration into injured muscle one day after a neonatal incision (Fig. 1B-C). Both vehicle and AP treated groups displayed significant paw guarding one day after a neonatal incision, with no statistical difference between groups (Fig. 1D, *p<0.001 vs. baseline (BL), p=0.071 n.inc+AP vs. n.inc+vehicle). However, while vehicle treated animals displayed a reduction in mechanical withdrawal thresholds to muscle squeezing one day after incision (*p=0.014 vs. BL), AP treated animals were hyposensitive compared to baseline and controls (Fig. 1E, *p=0.007 vs. BL, ^p=0.028 vs. n.inc+vehicle).

**Figure 1:**
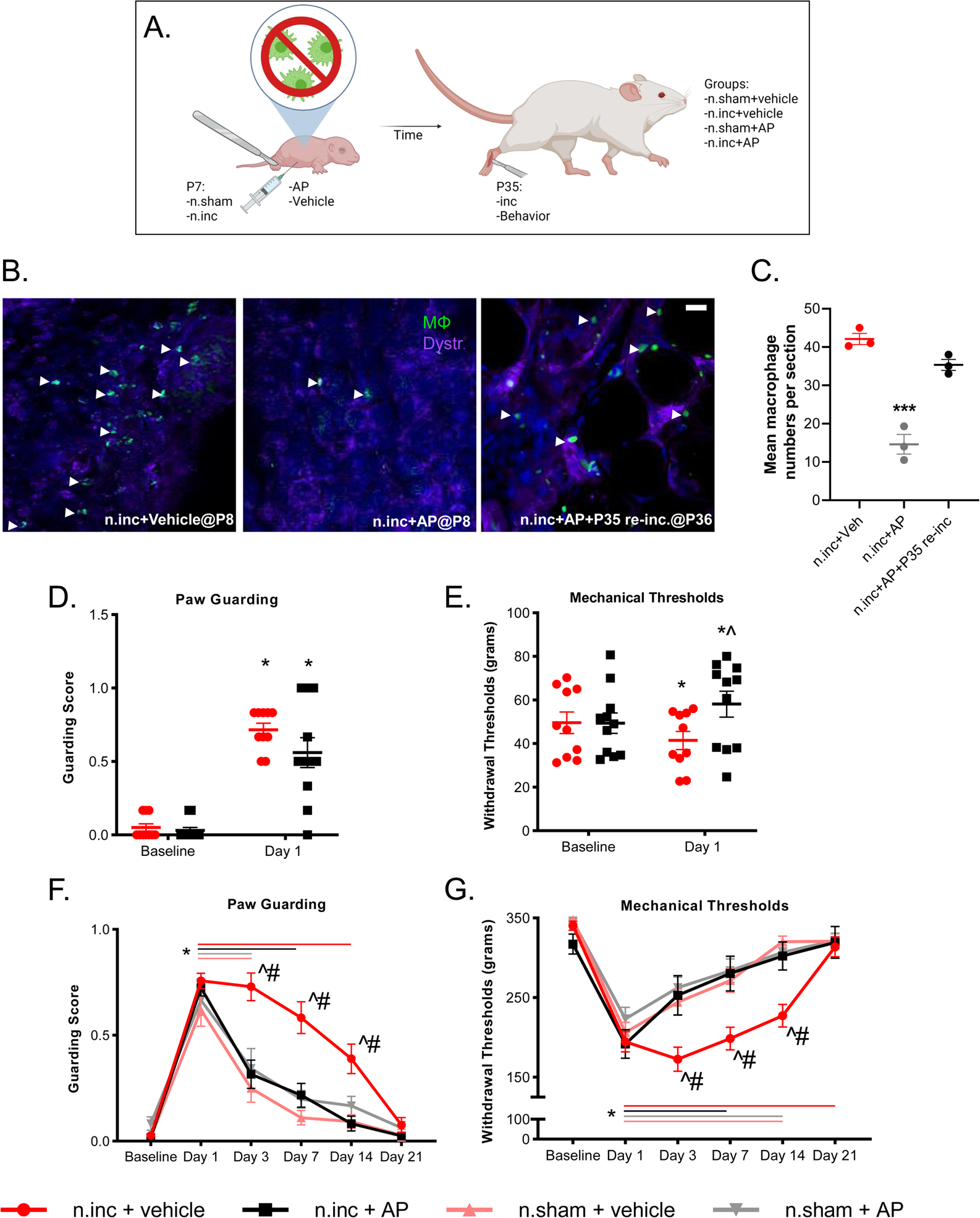
Early life ablation of macrophage alters responses to re-injury later in life. **A.** Schematic of the experimental design used in MaFIA mice. **B-C.** The treatment of MaFIA animals with AP early in life was sufficient to ablate macrophages at the injury site one day after the neonatal incision compared to vehicle treated controls. Macrophages are present in the hind paw of a P35 injured animal even after neonatal AP treatment (***p<0.001 vs. n.inc+vehicle and n.inc+AP+P35 re-inc.; one way ANOVA,Tukey’s; n=3 per group). Scale bar, 10μM. **D.** Paw guarding one day after a neonatal incision with or without macrophage ablation indicated an effect of day (F=103.418) but no difference between groups (*p<0.001 vs. BL for AP and vehicle treated animals; p=0.071 AP treated vs. vehicle treated). **E.** Muscle squeezing in the same animals showed an interaction for withdrawal thresholds (F=16.647). Vehicle treated animals had a significant reduction from baseline (BL) (*p=0.014 vs. BL) while incision+AP animals were hyposensitive and significantly different from controls (*p=0.007 vs. BL; groups were ^p=0.028 vs. controls; two-way RM ANOVA; n=10-11/group). **F.** Animals that received an early life ablation of macrophages (n.inc+AP) did not have altered baseline scores at adolescence, but the response to a second injury revealed a significant effect of dual incision (guarding F=5.635, muscle squeezing F=5.303). Groups that received a sham as neonates (n.sham+vehicle, n.sham+AP) guarded through day 3 (*p<0.01 vs. BL) while vehicle treated dually incised animals guarded through day 14 (*p<0.001 vs. BL). Animals from the neonatal incision (n.inc+AP) group returned to BL guarding scores after seven days (*p<0.05 vs. BL). At days three, seven, and 14, the n.inc+vehicle group was significantly different compared to all other groups (^p<0.05 n.inc+vehicle vs. controls, #p<0.001 vs. n.inc+AP). **G.** In the same group of animals, a similar pattern was observed for muscle squeezing withdrawal thresholds. N.sham operated groups were hypersensitive through day 7 (*p<0.005 vs. BL). The n.inc+vehicle group showed reduced thresholds through day 14 (*p<0.001 vs. BL), but n.inc+AP treated animals were hypersensitive only through day 3 (*p<0.001 vs. BL). At days three, seven and 14, the n.inc+vehicle group had significantly lower thresholds compared to all other groups (^p<0.005 vs controls, #p<0.005 vs. n.inc+AP). Colored horizontal lines indicate the duration of significance compared to BL for each group. Data shown as mean +/-SEM.

We then allowed different animals to age to P35 when sensory neurons and immunological functions are more developed^6,7,43,44^. At this time point, we first performed evaluations to determine any long-lasting effects of early life incision. Prior to a second incision at P35, we evaluated muscle integrity and LysM+ cell presence (Supp. Fig. 1C). Evans Blue Dye (EBD), when administered peripherally, is unable to penetrate into myofiber bundles unless they are damaged or undergoing repair^45^. Two days after adolescent injury, we observed that about half of the myofibers were positive for EBD. However, we found significantly less EBD staining within myofibers in animals that received an early life injury or early life sham (Supp. Fig. 1D-E, *p<0.001 vs. two days post incision). In addition, there were very few LysM+ cells still present in the muscle of monocyte/macrophage reporter animals (LysM;tdTom) at P35 after neonatal sham (n.sham) or incision surgery (n.inc) (Supp. Fig. 1F, p=0.648 vs. n.sham).

To determine the effect of early life injury in the presence or absence of macrophages on re-injury pain-like behaviors, we evaluated MaFIA animals treated with AP or vehicle early in life that received either a P7 sham or incision surgery. After aging and prior to a second incision, we observed no differences between any groups for paw guarding or muscle squeezing. One day after the P35 incision, we confirmed that even in neonatal AP treated animals, muscle *Csf1r+*, GFP expressing macrophages are present in the injured muscle (see Fig. 1B, C). At this same time point, we found expected acute pain-like behaviors from all groups. However, n.sham animals from both treatments returned to baseline levels of paw guarding after day three (Fig. 1F, *p<0.05 vs. BL for all groups) and muscle withdrawal thresholds after day seven (Fig. 1G, *p<0.001 vs. BL for all groups). We detected no effect of neonatal AP alone on nociceptive development in n.sham+AP animals (n.sham+vehicle vs. n.sham+AP). Unlike n.sham animals, n.inc+vehicle animals displayed long-lasting pain-like behaviors after a second incision through day 14 for guarding (Fig. 1F, *p<0.001 vs. BL, ^p<0.05 vs. n.sham+vehicle, #p<0.001 vs. n.inc.+AP) and day 21 for muscle squeezing (Fig. 1G, *p<0.001 vs. BL, ^p<0.005 vs. n.sham+vehicle, #p<0.005 vs. n.inc+AP). Early life ablation of macrophages rescued this persistent pain-like phenotype with n.inc+AP animals displaying guarding and mechanical hypersensitivity only through day seven (Fig. 1F-G, guarding *p<0.05 vs. BL; squeezing *p<0.001 vs. BL). Animals of this group were not significantly different at any time point compared to controls.

To evaluate if the ablation of *Csf1r*+ macrophages early in life alters the sensitization of sensory neurons following dual incision, we performed intracellular recordings of individual sensory neurons in an *ex vivo* intact muscle/tibial-nerve/DRG/SC preparation (Fig. 2A). We evaluated multiple peripheral response properties seven days after the second incision in vehicle and AP treated mice where we saw a robust block of both spontaneous guarding and evoked muscle squeezing behaviors (see Figs. 1F-G). We found no difference in the conduction velocity (CV) of thinly myelinated group III afferents in animals treated with AP versus vehicle seven days after the second incision (Fig. 2B, p=0.987). However, unmyelinated group IV afferents showed a significantly slower CV in AP treated animals versus vehicle treated animals (Fig. 2C, *p=0.048). After mechanical stimulation of receptive fields, we observed no change in average mechanical force necessary to evoke an action potential (Fig. 2D, p=0.430).

**Figure 2.**
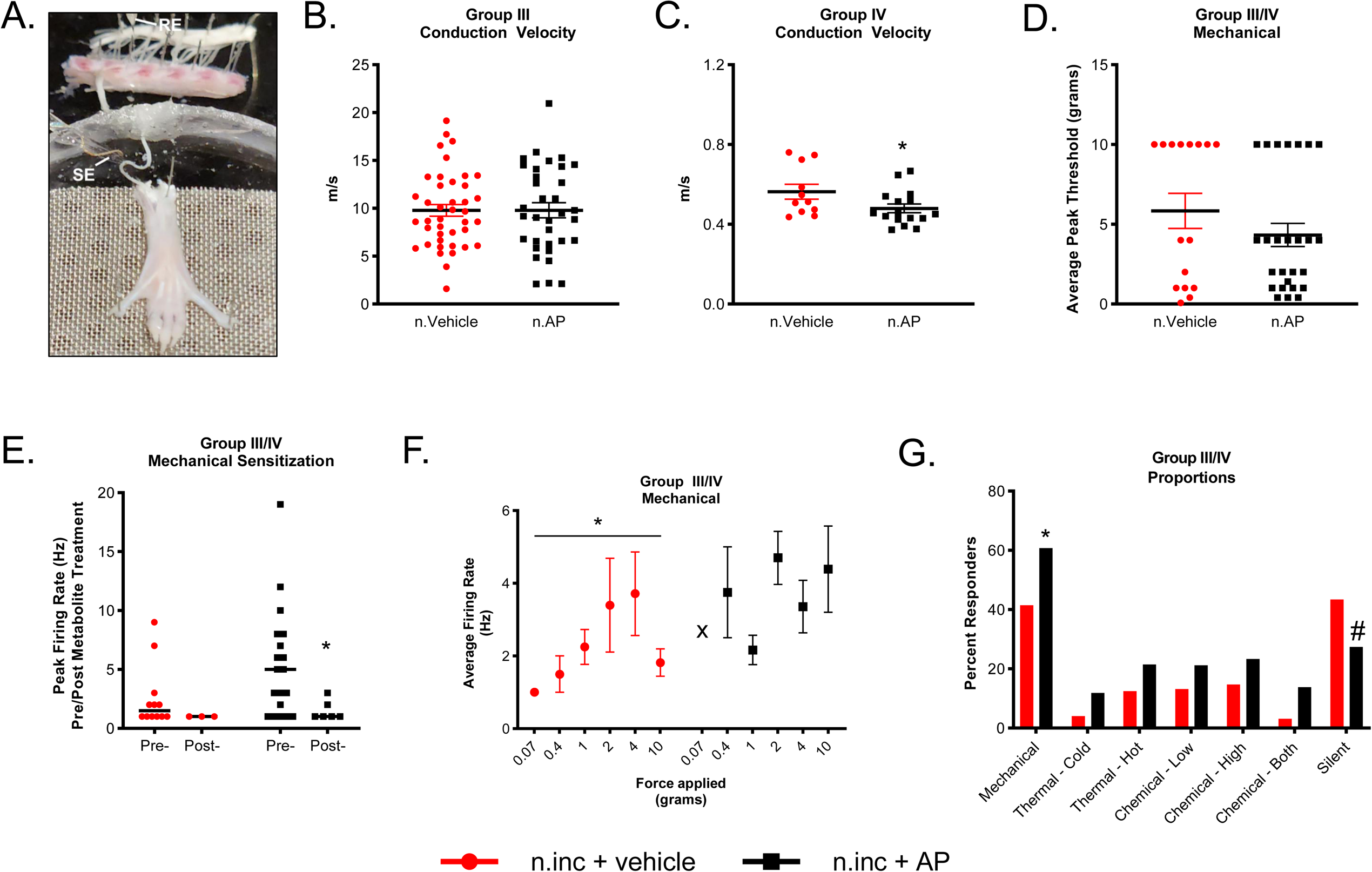
Macrophage depletion in neonates alters sensory neuron sensitization following repeated incision. **A.** Representative image of the intact electrophysiological preparation containing the injured hind paw muscle, tibial nerve, DRGs and SC. The search electrode (SE) and recording electrode (RE) are marked. **B.** There is no difference in CVs between groups in group III muscle afferents (p=0.987 n.inc+AP vs. n.inc+vehicle, n=34 and 39 respectively, one way ANOVA, Tukey’s) but (**C**) there is a significant reduction of group IV conduction velocities (H=3.896) in the neonatal treated AP group compared to vehicle (*p=0.048 n.inc+AP vs. n.inc+vehicle, n=16 and 11 respectively, ANOVA on Ranks, Dunn’s). **D.** There are no differences in overall mechanical thresholds between groups (p=0.430 n.inc+AP vs. n.inc+vehicle, n=31 and 22 respectively, ANOVA on Ranks, Dunn’s). **E.** There is no change in mechanical firing rate pre-to post-metabolite treatment in vehicle treatment group, unlike that seen in the AP treatment group (H=4.934, n.inc+vehicle: p=0.223, n.inc+AP: *p=0.031, ANOVA on Ranks, Dunn’s). **F.** There is a significant increase in firing rate over mechanical forces applied in the vehicle treatment (H=5.760) but not in the AP treatment group (n.inc+vehicle: *p=0.019, One way ANOVA on Ranks, Dunn’s. n.inc+AP: p=0.149, One way ANOVA on Ranks, Dunn’s, “X” indicates no cell responded to the applied force). **G.** There is an increase in the number of mechanically sensitive cells in AP treated animals compared to vehicle treated animals (*p=0.049 n.inc+AP vs. n.inc+vehicle, Chi-Square on mechanical responders), with a corresponding non-significant decrease in cells that responded to no stimulus (“silent”, #p=0.090 n.inc+AP vs. n.inc+vehicle, Chi-Square on silent neurons). Data shown as mean +/-SEM.

Overall, we observed no difference in the firing dynamics on all neurons collectively (Supp. Fig. 2), but upon the analysis of subtypes, we found that chemosensitive and mechanically sensitive fibers (without cold responsiveness^46^), had an overall greater mechanical firing rate that was reduced following incubation with noxious metabolites (Fig. 2E, *p=0.031). Vehicle controls did not show a pre/post-metabolite difference. We further uncovered that control nociceptors displayed a direct increase in firing rate as we applied greater forces to the receptive field as has been previously reported^47^. However, in animals with an early in life restricted knockout of macrophages, we found no association between firing rate and force applied (Fig. 2F, *p=0.019, main effect within n.inc+vehicle). We found no differences in thermal or chemical firing rate or instantaneous frequencies between groups (Supp. Fig. 2). The proportion of responders in AP treated animals resulted in a significantly higher number of responders to mechanical stimuli (42% increased to 61%) with a corresponding, non-significant, reduction (43% decreased to 27%) in the number of non-responder “silent” neurons (Fig. 2G, *p=0.049, #p=0.090) compared to vehicle controls. We observed no change in the proportion of polymodal neurons (Supp. Fig. 2). Together, these data indicate that the temporary loss of macrophages during neonatal injury changes inherent sensory neuron dynamics and decreases the ability of nociceptors to change their responsiveness over increasing mechanical forces.

### Adoptive transfer of “primed” macrophages drives increased mechanical hypersensitivity

We next wanted to determine if macrophages isolated from neonatal incised animals were sufficient to evoke a ‘primed’ response in older mice. To do this we performed adoptive transfer (AT) experiments in which macrophages from one animal (donor) were isolated and transferred to a sex and age-matched naïve host (Fig. 3A). The *Csf1r* driver in MaFIA animals also promotes GFP expression so we used this strain to isolate and sort peritoneal GFP+ macrophages (Fig. 3B). We transferred the isolated cells into to the right hind paw of naïve hosts at P35. We first ensured that the AT was accepted by dissecting the site of injection as well as other immune-related tissues. We found GFP+ cells in WT hosts three weeks after the AT specifically at the muscle injection site and not in the nearby bone marrow, lymph node, nor spleen (Fig. 3C). After confirming the transfer, we found that AT of unprimed macrophages by itself (n.sham+AT alone) into an uninjured paw had no effect on paw guarding, but injection of primed cells (n.inc.+AT alone) by itself induced low-level paw guarding that lasted for one day (Fig. 3D, *p=0.044 vs. BL), and slightly reduced muscle squeezing thresholds, with no difference between groups (Fig. 3E, *p<0.05 Day 1, Day 3 vs. BL). To determine how exposure to injury might evoke a differential behavioral response, we also paired the AT with a P35 incision in different animals. We found that primed cells (n.inc.+AT with P35 inc), but not unprimed cells (n.sham+AT with P35 inc), when transferred into an injury environment increased mechanical hypersensitivity compared to controls measured by muscle squeezing (Fig. 3E, ^p<0.01 Day 1, Day 3 vs. n.sham+AT alone). This effect was not observed in paw guarding where donor neonatal injury did not impact host behavioral outcomes, indicating a modality-directed phenotype. These data indicate that an early life injury drives changes to macrophages which are sufficient to drive some of the enhanced behavioral phenotypes due to injury in a host environment.

**Figure 3:**
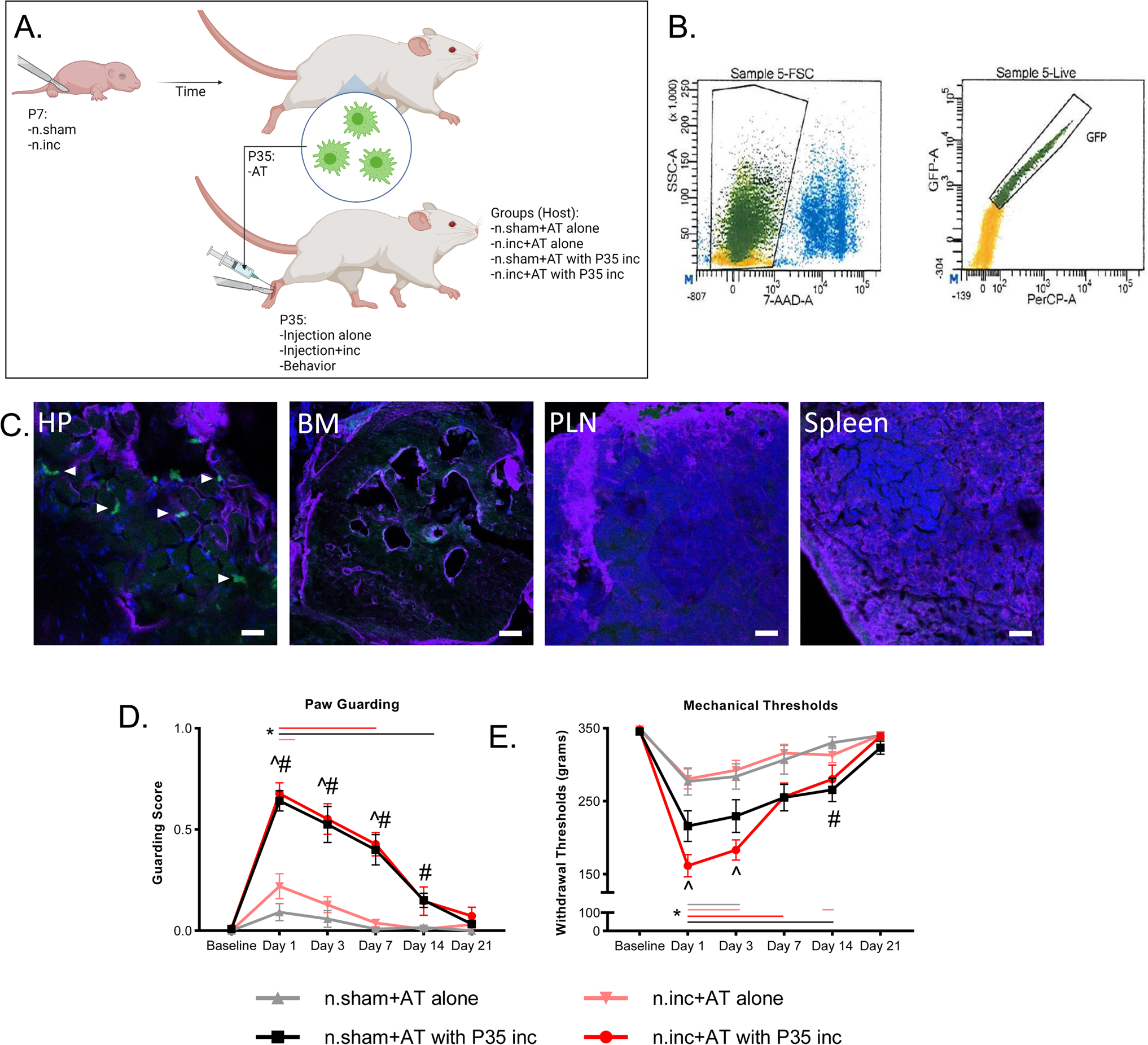
Adoptive transfer of macrophages from neonatal incised animals increases pain-like behaviors. **A.** Outline of the experiment design for AT experiments. **B.** Example of the sorting of live GFP+ peritoneal macrophages from MaFIA mice. **C.** Example images demonstrating the presence of GFP+ donated cells (arrows) in the hindpaw of a host 21 days after injection, but not in the bone marrow (ipsilateral tibia), lymph node (potineal), or spleen. Scale bars, 25μM; BM, 150μM. **D.** For guarding behaviors, there was an effect (F=3.132) of macrophage exposure to injury. Both primed (n.inc+AT with P35 inc) and unprimed (n.sham+AT with P35 inc) cells injected into an incision site induced prolonged guarding (*p<0.05 vs. BL). In the absence of injury at P35, primed (n.inc+AT alone) cells induced one day of guarding (*p=0.044 vs. BL) while unprimed cells (n.sham+AT alone) did not induce detectable behavior. There was a significant difference between n.inc+AT with P35 inc and n.sham+AT alone at days 1, 3 and 7 (^p<0.001 vs. n.sham+AT alone) and between n.sham+AT with P35 inc vs. n.sham+AT alone at days 1-14 (#p<0.05 vs. n.sham). **E.** The same animals were then tested for muscle withdrawal thresholds by muscle squeezing which resulted in a significant interaction (F=4.318). All groups had a significant reduction in withdrawal thresholds that varied over time depending on condition (*p<0.05 vs. BL). N.inc+AT with P35 incision was significantly different than n.sham+AT alone at days 1 and 3 (^p<0.01 vs. n.sham+AT alone). N.sham+AT with P35 incision vs. n.sham+AT alone treatment showed an effect at day 14 (#p=0.015 vs. n.sham+AT alone). Two-way RM ANOVA, Tukey’s post hoc, n=8-11/group. Colored horizontal lines indicate the duration of significance compared to BL for each group. Data shown as mean +/-SEM.

### Early life incision alters the peripheral macrophage epigenetic landscape

To determine what molecular changes might be occurring in these immune cells because of early life incision, we collected peritoneal macrophages from three different conditions of reporter animals (LysM;tdTom): naïve animals with macrophages isolated at P7 (naïve isolated at P7), naïve macrophages isolated at P35, and P7 incised animals with macrophages isolated at P35 (P7 inc, isolated at P35 (primed)) (Fig. 4A). To test if early life injury altered the epigenetic landscape of macrophages, we performed an Assay for Transposase-Accessible Chromatin sequencing (ATAC-seq) to measure chromatin accessibility. Quality control confirmed our isolation and predicted that the sequenced cells were macrophages (Supp. Fig. 3). Principle component analysis (PCA) revealed distinct clustering of the three groups, indicating an overall effect of early life incision (Fig. 4B). We repeated these same conditions to evaluate gene expression using RNA-seq and found somewhat similar clustering in our bulk RNA-seq PCA (Fig. 4C). We evaluated the top 100 differentially accessible chromatin sites (Fig. 4D) and top 100 differentially expressed genes (DEG) (Fig. 4E) from age-matched macrophages in both the P35 naïve and primed groups. We obtained 10 candidate genes that were both differentially accessible and expressed in the same direction because of early life incision (Fig. 4F). From this sequencing data, we found that a promising candidate, NGFR (aka. p75NTR), was differentially regulated in macrophages due to neonatal injury (Fig. 4G-H). p75NTR has long been of interest for nervous system development and nociception, and it has roles in macrophage differentiation and function^28,31^. We therefore decided to test whether macrophage p75NTR could modulate neonatal nociceptive priming phenotypes.

**Figure 4.**
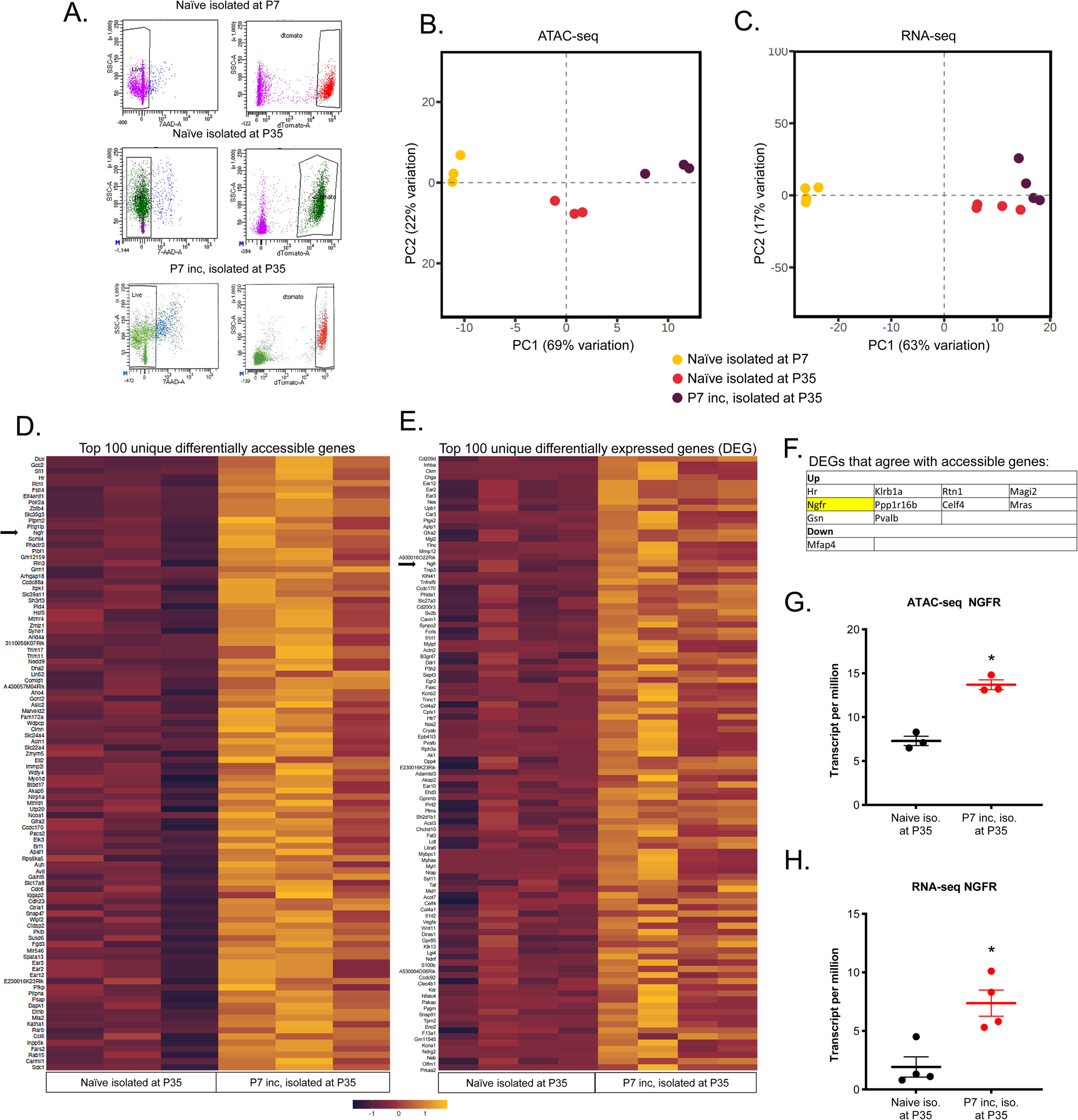
Early life incision alters the epigenetic and mRNA landscape of LysM+ cells. **A.** Example sorting of peritoneal macrophages from P7 naïve, P35 naïve, and P7 incised animals isolated at P35. **B.** Principle component analysis from ATAC-seq datasets demonstrating distinct clustering of groups. **C.** Principle component analysis form the RNA-seq dataset shows similar distinct clustering of groups. **D.** The top 100 differentially accessible chromatin regions in macrophages in P35 naïve cells compared to P35 primed cells (age matched). **E.** The top 100 differentially expressed genes (DEGs) for the same conditions. **F.** A table of factors that had differentially accessible regions and DEGs. NGFR (p75NTR) is indicated. **G.** ATAC-seq indicating higher reads of NGFR (*p=0.001 vs. controls, one-way ANOVA, Tukey’s, n=3-4/group), and (**H**) transcripts per million (TPM) data for RNA-seq (*p=0.008 vs. controls). Data shown as mean +/-SEM.

To do this, we crossed the inducible LysMCreERT2 animal with a p75NTR floxed animal^48^. All littermates were first incised at P7 to induce priming. Beginning at P33 and continuing until P37, animals were injected with tamoxifen to eliminate p75NTR from LysM+ cells. At P35, all animals received a second incision to evoke the primed macrophage and behavioral responses. Behavior was evaluated prior to and for three weeks following the P35 incision (Fig. 5A). To verify our injection protocol, we obtained samples following the final injection of tamoxifen at P37. We found an overall decrease of p75NTR mRNA from total peritoneal isolated cells in knockout animals compared to controls (–43±51% LysMCreERT2;p75fl/fl vs. controls). A P35 incision resulted in a number of F4/80+ macrophages within the injured tissue, of which 31% were positive for p75NTR in control genotypes (Fig. 5B). The knockout of p75NTR from LysM+ cells resulted in a significant reduction in co-stained (F4/80+ and p75+) macrophages (Fig. 5C, *p<0.05 vs. controls). Behaviorally, we observed that the knockout of p75NTR specifically in macrophages did not prevent long-lasting paw guarding (Fig. 5D), but significantly blunted the severity of the mechanical sensitivity one day following injury compared to controls (Fig. 5E; ^p=0.01 vs. controls). We further determined that hypersensitivity of the contralateral paw (Supp. Fig. 3, contralateral data from Fig. 1) was also inhibited by p75NTR knockout in LysM+ cells (Supp. Fig. 3). These data indicate that p75NTR has a partial but critical role in the macrophage-driven cellular memory that drives nociceptive priming.

**Figure 5.**
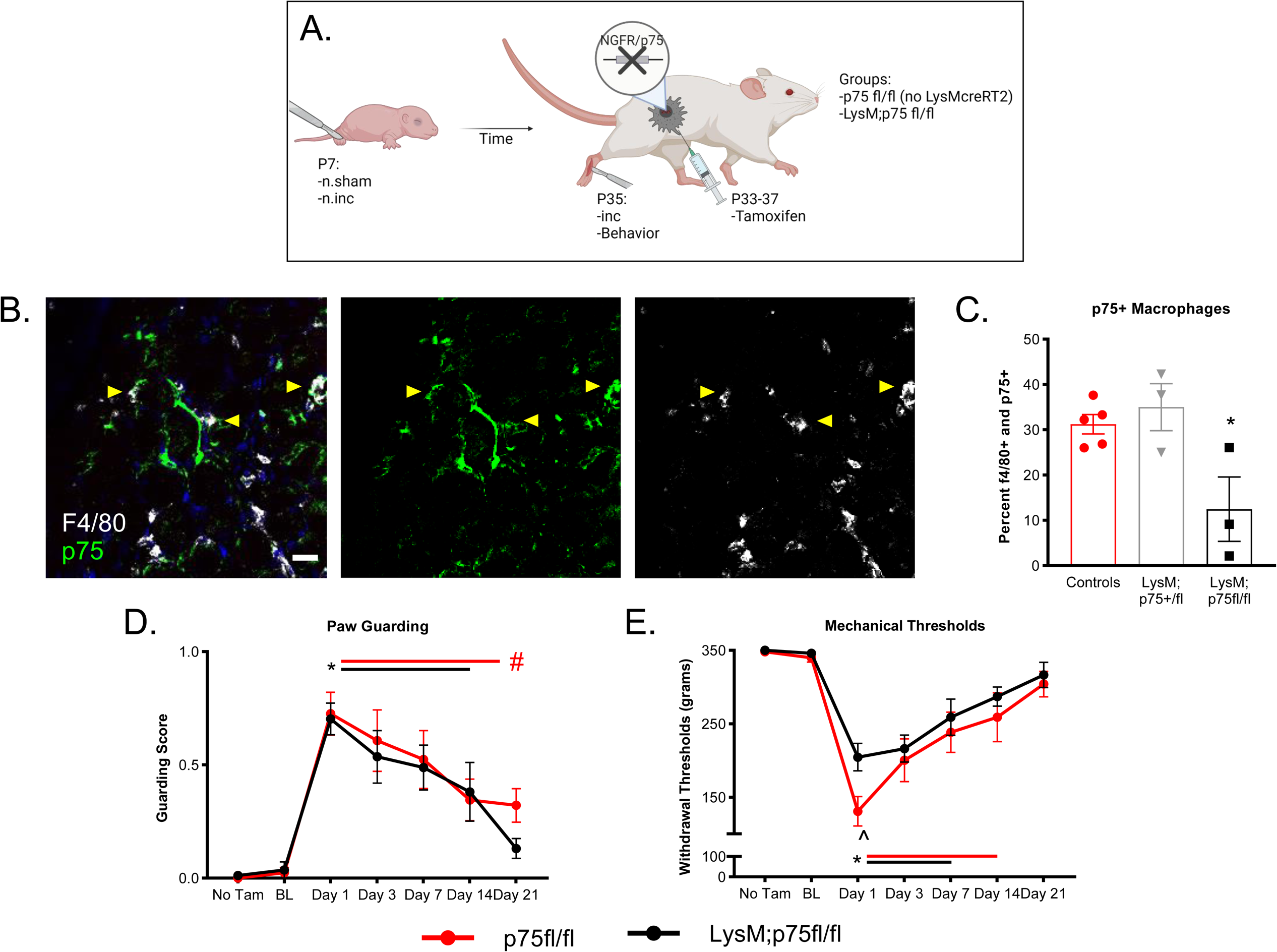
Macrophage specific knockout of p75NTR after neonatal incision impacts neonatal nociceptive priming. **A.** Schematic of knocking out p75NTR from macrophages following neonatal incision. **B.** Representative images of p75NTR labeling (green) co-stained with F4/80 (macrophages, white) in the injured hind paw. Yellow arrows indicate double positive staining. Scale Bar, 25μM. **C.** There is a significant reduction (F=4.012) in the percentage of double positive cells in LysM;p75fl/fl animals compared to other genotypes (*p<0.05 vs. controls (p75+/fl and p75fl/fl) and LysM;p75+/fl, one way ANOVA, Tukey’s, n=3-5/group). **D.** We found no effect of tamoxifen prior to the second injury (No Tam vs. BL time points) on paw guarding. Following the second incision there was robust paw guarding (F=26.258), with a faster recovery in the LysM;p75NTRfl/fl mice (*p<0.05 vs. BL, #p=0.05 vs BL p75fl/fl, Two-way RM ANOVA, Tukey’s). **E.** No effect of tamoxifen on mechanical withdrawal thresholds was found prior to injury (No Tam vs. BL time points), however there was an effect following a second incision (F=40.516). Both groups displayed mechanical sensitivity on days one, three and seven (*p<0.01 vs. BL). The p75fl/fl genotype continued to display significant sensitivities at day 14 while LysM;p75fl/fl animals did not (p75fl/fl: *p=0.002 vs. BL, LysM;p75fl/fl: p=0.063 vs. BL). At one day following second incision, LysM;p75fl/fl animals were less hypersensitive than controls (^p=0.01 vs. controls). Two-way RM ANOVA, Tukey’s, n=7/group.). Colored horizontal lines indicate the duration of significance compared to BL for each group. Data shown as mean +/-SEM.

### Knockout of p75NTR in macrophages alters neuroimmune communication in mouse and in human iPSCs

We then wanted to test if macrophage p75NTR modulated nociceptive priming through its role in inflammation in a system that could maintain a long-term immune memory. To do this, we began evaluating bone marrow derived macrophages (BMDMs) and how they respond to neurotrophins. First, we isolated bone marrow from naïve animals at P35, differentiated them into macrophages^49^, and measured cytokine release in response to different stimuli using cytokine arrays. We tested the effect of knocking down p75NTR using an siRNA strategy (example arrays Fig. 6A; siRNA transfection Supp. Fig. 4). We tested the effect of stimulation of two neutrophins, NGF and BDNF, which both bind to p75NTR in addition to TrkA and TrkB, respectively. We compared these to negative (vehicle) and positive (lipopolysaccharide and interferon gamma (LPS+IFNγ)) controls (Fig. 6B; full data Supp. Fig. 4). In murine cells, the knockdown of p75NTR in macrophages alone (sip75+Vehicle (Veh)) affected both pro-and anti-inflammatory cytokines. If stimulated with LPS+IFNγ, the response was predominately pro-inflammatory. NGF did not appear to alter many proteins alone in control cultures (Supp. Fig. 4), however, NGF+p75NTR knockdown caused a significant down regulation of pro-inflammatory cytokine release and increased anti-inflammatory cytokine production (*p<0.05 vs. siCON+Veh, ^p<0.05 vs. sip75+LPS+IFNγ, #p=0.05 vs. sip75+Veh, ##p>0.05 vs siP75+Veh; Fig. 6C). Fewer changes were observed when BDNF or a combination of NGF and BDNF was used (Supp. Fig. 4). Together with our previous data, this may indicate that the epigenetic regulation of p75NTR in macrophages that are exposed to early life injury could promote pro-inflammatory/pro-nociceptive factor release and the targeting of this receptor can reverse this profile.

**Figure 6.**
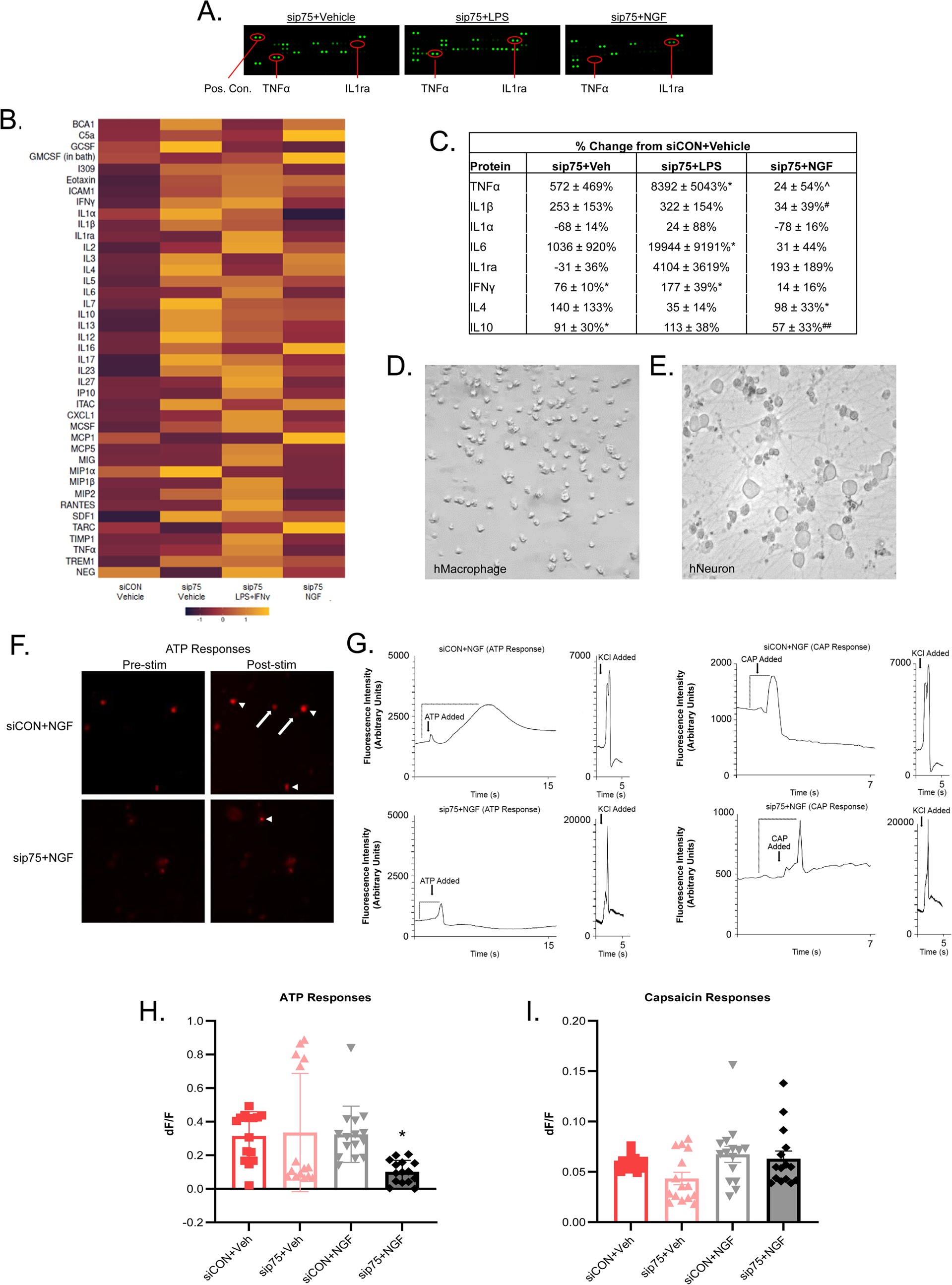
p75NTR expression in macrophages alters the inflammatory signature of macrophages and neuronal activity. **A.** Examples of the protein arrays that were run on the lysates of BMDMs that received a knockdown of p75NTR and were stimulated with indicated factors. **B.** Heatmap indicating the protein expression detected across each stimulus, n=4/group. **C.** Examples of significantly regulated proteins detected from the protein array (*p<0.05 vs. siCON+Veh, ^p<0.05 vs. sip75+LPS, #p=0.05 vs. sip75+Veh, ##p>0.05 vs sip75+Veh; one-way ANOVA, Tukey’s or ANOVA on Ranks, Dunn’s. n=4/group). **D-F.** Representative images of iPSC-derived macrophages, sensory neurons and Rhod-2 responses to ATP. Arrows indicate increased responses. Arrowheads indicated new responses. **G.** Example calcium transients from iPSC derived sensory neurons in response to ATP, capsaicin or KCl stimulation. The arrow indicates when the ATP or capsaicin was added, and the brackets indicate the part of the trace that was analyzed to calculate changes in fluorescence intensity in direct response to a stimulus. Scale Bar, 20μM. **H.** Exposure of iPSC derived sensory neurons to media from macrophages with loss of p75NTR treated with NGF caused decreased responsiveness of neurons to ATP (F=4.257) (*p<0.04 vs. all other conditions, one way ANOVA with Tukey’s post hoc, n=15 cells per group). **I**. No changes in calcium responses to capsaicin (F=2.702) were observed (one way ANOVA). Data shown as mean +/-SEM or percent change from controls with variance.

To test if these effects are conserved in human cells to influence sensory function, we then performed similar experiments with NGF and p75NTR knockdown in human iPSC-derived macrophages (Fig. 6D). We extracted the media of vehicle or NGF stimulated iPSC-derived macrophages that were treated with either control or p75 targeting siRNAs and applied it directly to human iPSC-derived sensory neurons (Fig. 6E) loaded with the calcium indicator Rhodamine-2 (Rhod-2; Fig. 6F). After 24 hour incubation, we discovered no difference between control groups that received conditioned media from macrophages with NGF or vehicle, but we did find a decreased calcium response to ATP in iPSC derived sensory neurons that were previously incubated with conditioned media from human macrophages treated with sip75+NGF (Fig. 6G-H). No differences between groups were found after treatment with capsaicin (Fig. 6I). Analyzed cells were confirmed to be viable and healthy by responding to high KCl stimuli (Fig. 6G inserts). Together with our previous data, this suggests that neuroimmune signaling in murine and human cells is impacted by p75 expression.

### Transplant of neonatal “primed” bone marrow alters female incision-related behaviors

Trained immunity describes the concept of a long-lasting cellular memory that allows the immune system to display an enhanced response to subsequent insults. To determine the persistence of neonatal priming *in vivo*, we repeated priming experiments from Figure 1. However, in these experiments, we waited 20 weeks following neonatal incision, when animals were P147, to perform the repeated HP incision followed by three weeks of behavioral testing. We found a persistent effect of neonatal priming in a sex specific manner. While the effect of neonatal incision was lost in the aged males (Fig. 7A, B), re-injured females displayed long-lasting spontaneous paw guarding (Fig. 7C ^p=0.02 n.inc+P147 inc female vs. n.sham+P147 female at day 7) and mechanical hypersensitivity (Fig. 7D ^p<0.01 n.inc+p147 inc female vs. n.sham+P147 female at days 7 and 21) compared to neonatal females that received a sham surgery.

**Figure 7.**
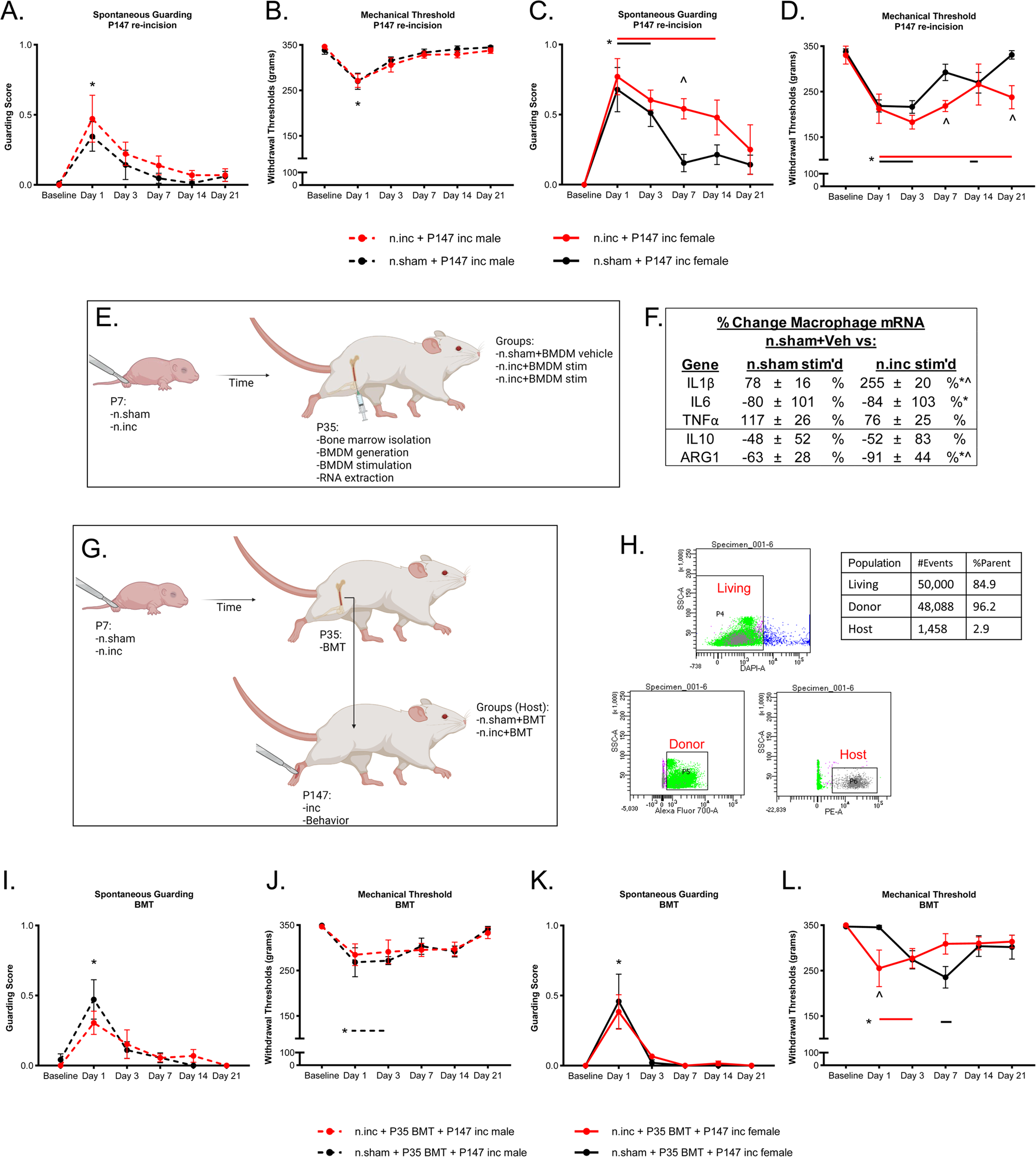
Bone marrow transplant from neonatal incised animals transfers a “pain memory” to naïve animals. **A, C.** There is a main effect of group (F=6.96) and day (F=27.20) for guarding with post hoc analysis revealing that n.inc+P147 inc females guard through day 14 (*p<0.05 vs. BL) and significantly more than n.sham +P147 females at day 7 (^p=0.02). n.sham+P147 females guard through day 3 (*p<0.001 vs. BL) while both male groups only guard at day one (*p<0.05 vs. BL). Two-way RM ANOVA, Tukey’s. **B, D.** In the same animals for muscle squeezing there is an interaction effect (F=3.50). n.inc+P147 inc female have significantly reduced thresholds from baseline for three weeks (*p<0.05 vs. BL) and are significantly more hypersensitive than n.sham+P147 inc females at days seven and 21 (^p<01 vs. n.sham+P147 inc female). N.sham+P147 inc females have reduced thresholds at days one, three and 14, while males from both groups are hypersensitive at day one only (*p<0.05 vs. BL). Two-way RM ANOVA, Tukey’s, n=11-13/injury condition, 4-7/sex in condition. **E.** A schematic indicating timeline for neonatal nociceptive priming experiments separating injury by 140 days. **F.** Analyses of the transcripts from BMDMs after stimulation with LPS+IFNγ (*p<0.05 vs. n.sham no stimulation controls, ^p<0.05 vs. n.sham with stimulation (stim’d), n=3-8/group, Two-way ANOVA, Tukey’s). **G.** Schematic of the bone marrow transplant experiments. **H.** Representative cell sorting of living cells, 45.1+ host remaining cells and 45.2+ transferred cells. **I, K.** After recovery from the BMT, there is a main effect of day (F=22.90). Animals display no guarding behaviors prior to incision. Following an incision both n.sham and n.inc groups from both sexes guarded for one day only (p<0.01 vs. BL, Two-way RM ANOVA, Tukey’s). **J, L.** In the muscle squeezing assay, n.inc+P35 BMT+P147 inc. females are hypersensitive to baseline at days one and three (*p<0.05 vs. BL) and have significantly lower thresholds than controls at day one (^p=0.013 vs. n.sham+P35 BMT+P147 inc female). n.sham+P35 BMT+P147 inc females do not begin to display sensitivity until day seven (*p=0.001 vs. BL). N.sham+P35 BMT+P147 inc males have reduced thresholds from BL at days one and three (*p<0.05 vs. BL). Two-way RM ANOVA, Tukey’s, n=10-11/injury condition, 4-6/sex in condition. Colored horizontal lines indicate the duration of significance compared to BL for each group. Data shown as mean +/-SEM.

We next hypothesized that underlying changes to bone marrow cells caused by an early life injury could be a mechanism of this retained behavioral and molecular “memory.” Therefore, we isolated mouse BMDMs at P35 from neonatal incised or sham controls. We found no difference in the total number of bone marrow cells isolated due to an incision (Supp. Fig. 4). However, when we stimulated these cultured macrophages with LPS+IFNγ as an artificial second “injury” (Fig. 7E) we observed that early life injury “primed” these cells as they responded with a greater pro-inflammatory profile compared to n.sham controls (Fig. 7F, *p<0.05 vs. n.sham+vehicle, ^p<0.05 vs. n.sham+LPS+IFNγ (stim’d)). These results indicate that incision induces a long-lasting (at least 28 days) reprogramming of hematopoietic progenitors to generate macrophages that are primed to respond more potently to inflammation.

Finally, to test if hematopoietic stem cells (HSCs) are a mechanism through which this long-term memory is maintained and functionally relevant to nociception, particularly in females, we performed a bone marrow transplant (BMT) experiment. To do this, we performed a neonatal sham or incision, allowed the animals to age until P35 (where we detected an altered BMDM expression pattern), and then performed a BMT into naïve adolescent mice (Fig. 7G). To regain full immune functional activity following a BMT, animals were allowed to recover for 16 weeks (P147) when almost all hematopoietic cells are derived from the donated hematopoietic stem cells^50,51^. At this time point animals were subjected to a hind paw incision and were monitored for another three weeks. We successfully transferred the bone marrow evidenced by the vast majority of CD45.2+ donor cells and a minority of CD45.1+ host cells (Fig. 7H). Prior to incision later in life, we detected an expected, but non-significant, increase in p75NTR transcript from bulk isolated peritoneal cells (128 ± 28%, One way ANOVA n=4-5/group). In this assay we could not isolate macrophages specifically so other immune cells such as B and T cells are contributing to these values and likely diluting the effect. Interestingly, we found that if we analyze female and male peritoneal cells separately, regardless of condition, females had 480 ± 25% more p75NTR than males (p=0.016, ANOVA on Ranks, Dunn’s, n=4-5/sex). When we evaluated all animal behavior independent of sex following later in life incision, we uncovered little evidence of enhanced pain-like behaviors in either group with quick recovery in both paw guarding and muscle squeezing (Supp. Fig. 5). Both groups displayed guarding behavior for only one day (p<0.001 vs. BL) and variable mechanical hypersensitivity for 1-7 days (p<0.05 vs. BL; Supp. Fig. 5). However, when we separated animals by sex, although the length of injury-induced behaviors was not different between groups and there was no acceleration or delayed onset sensitivity in males (Fig. 7I-J), we found a delayed onset of n.sham+BMT female-specific mechanical sensitivity. However, the n.inc+BMT (primed) females displayed a leftward shift from controls (Fig. 7K-L). One day after injury, female n.sham+BMT controls were not mechanically sensitive but female n.inc+BMT primed-cell recipient females had significantly lower thresholds than controls (^p=0.013 vs. controls). Together, these data describe an effect of early life incision on HSCs that can influence evoked pain-like behaviors for over 100 days in females. Therefore, the predisposition to chronic pain due to neonatal priming is enhanced in a sex-specific manner partially due to an HSC-related immune memory of early life injury.

## Discussion

In this study, we uncovered a mechanism in which the peripheral immune system can form and maintain a “pain memory” following a neonatal hind paw incision to modulate prolonged responses to repeated injury (graphical abstract). Macrophages were found to be both necessary (Figs. 1-2) and partially sufficient (Fig. 3) to drive neonatal nociceptive priming responses at the behavioral and afferent levels. This single, local insult dramatically altered the systemic macrophage epigenetic/transcriptomic landscape (Fig. 4) and function in part through the p75NTR receptor in mouse and human cells (Figs. 5-6). The molecular effects of early life injury were also evident in BMDMs, and following a BMT, which induced a sex-specific alteration in pain like behavior that was functionally relevant long-term in hosts (Fig. 7). This study is the first to show a macrophage cellular memory formed in neonates that modulates pain related outcomes. Together, these data help us to understand how the distinct cellular systems are changed by an aversive event in early development to influence nociceptive processing across the lifespan.

Previous work has evaluated the vulnerable periods that exist in the somatosensory and immune systems^5,38,52–57^ as well as genetic risk factors in children following surgery^32,58^. After neonatal injury, the inhibitory circuits and microglial activity of the spinal cord dorsal horn (SCDH) are dysregulated leading to hyperactivity of spinal output^59–64^. Moriarty et al found that a peripheral nerve block was sufficient to prevent neonatal nociceptive priming, while systemic morphine injection was not^13^. We previously demonstrated that peripheral growth hormone, which is a vital player in neonatal nociceptive development ^65^, can be used to prevent neonatal nociceptive priming ^5^. This highlights the importance of peripheral input to induce nociceptive priming and our work extends these findings to capture a neuroimmune mechanism.

Following an injury there are both local and global responses in the immune and nervous systems^38^. Macrophages are one of the first immune cells at the injury site to promote inflammation and subsequent wound healing. This is accomplished through damage associated molecular patterns (DAMPs) which can be stimulated by, and also control, cytokine and growth factor responses. Both DAMPs and the effector proteins released may also be pro-nociceptive such as high-mobility group box 1 (HMGB1), IL1β and NGF^66–71^. ATP is another DAMP that is present in our *ex vivo* preparation metabolite mixture which we found to alter the mechanical firing rate of chemosensitive neurons in animals that lacked macrophages during an early life incision (Fig. 2^67^). It may be that desensitization to a high metabolic/noxious tissue environment was a contributor to the rescuing of priming by early life macrophage ablation (Fig. 1). The conduction velocity slowing of group IV sensory neurons because of macrophage ablation is an interesting result. There are reports of activity dependent slowing of cutaneous unmyelinated fibers^72^, but this is unlikely to be the driving factor here because of the lack of spontaneous activity in these cells. There may be differences in the size of the population of neurons that could explain this phenomenon and could correlate with the differences in proportions of each type of fiber response detected (Fig. 2). AT experiments (Fig. 3) showed that macrophages are only partially sufficient to induce priming. This could suggest that other cells, including other infiltrating or resident immune cells, “nociceptive” Schwann cells^73^, or even sensory neurons themselves, may contribute neonatal nociceptive priming. However, these concepts would need further investigation.

Neonatal macrophages are unique compared to the adult, with a distinct epigenetic landscape (Fig. 4) and differential responses to stimuli^14,52,9,10,24,74^. The plasticity of the macrophage epigenome is critical to differentiate monocytes to macrophages, and acute stimulation of these cells induces epigenetic modifications^75–78^. Changes in macrophages at the epigenetic level is one way in which these cells become either “trained” or “tolerant” to subsequent stimuli^25,26^ ^57^. Recent work by Cobo et al reveals that infant nociceptive reflexes to noxious and non-noxious stimuli are altered by current or previous infection^79^. It is possible that an early life aversive event such as a surgical incision (or an infection) may generate molecular patterns that induce epigenetic priming in macrophages as was observed in the current study. Strategies to interrupt neuroimmune signaling may prove beneficial for these adverse injury-related effects^80–82^. It will be necessary to determine the factors that epigenetically train macrophages for future drug targeting.

While the priming initiation mechanism is yet unknown, we determined that p75NTR is one critical endpoint of the epigenetic modifications. In early development, neurotrophin signaling is necessary for sensory neuron specialization into distinct populations^83–86^, which can be modulated by p75NTR expression^31^. Macrophage p75NTR has been less studied, but HSCs express high levels of neutrophins, and we and others found that the inflammatory response from macrophages is directly affected by a loss of p75NTR (Fig. 6^29,87,88^). Developed immune cells regularly contain low levels of neurotropic factor receptors, but during inflammation they are robustly, and transiently, increased^5,28^. NGF in human derived macrophages modulates p75NTR expression and drives its co-localization with TrkA, which we found to have persistent upregulation in macrophages due to early life incision (data availability at NCBI: GSE224209)^88^. Further, Khodorova et al demonstrated that intraplantar NGF injections evoke expected pain-like behaviors, but that a general inhibition of the p75NTR receptor rescued the phenotype^89^. Combined with our data, this indicates that immune cell p75NTR increases their pro-inflammatory signature and that the ablation of this receptor can modulate pro-inflammatory signaling, blunt the ability of macrophages to communicate to sensory neurons, and partially prevent prolonged pain-like behaviors (Figs. 5-6).

Circulating immune cells that infiltrate injury sites are constantly replenished through the hematopoietic stem cell pool in the bone marrow. These are highly plastic cells that are differentially altered by injury type^90^, infection^91^ and development^28,92^. Our data support the notion that peripheral injury can alter these stem cells long-term, and changes in these cells are part of a system to maintain immune-related neonatal nociceptive priming (Figs. 4-7), however more work will be necessary to fully understand these findings. The mechanism in which HSCs are notified of noxious stimuli is currently unknown, although there are potential avenues to be tested including direct immune or neuronal signaling^93^, or indirect signals in the blood or cerebral spinal fluid^94^. Regardless, the results we find are reminiscent of trained immunity where an initial event is epigenetically retained in innate immune cells to enhance responses to subsequent activation^26^. In the context of immunity, the outcome is increased inflammation, while here the outcome was enhanced nociception. Evaluation of the functional effects of injury on HSCs by BMT indicated a female-specific hastening of mechanical hypersensitivity. Interestingly, we detected no effect in spontaneous paw guarding between early life incised and control animals indicating a modality specific effect (Fig. 7). It may be that “primed” macrophages have direct signaling roles that involve mechanical sensation as we saw similar, mechanically directed, effects with AT (Fig. 3) and p75NTR knockout (Fig. 5) behaviors. An understanding of how stem cells may be changed by peripheral injury may have important clinical considerations for a number of conditions that impact childhood development.

There are multiple technical points to consider. The depletion of macrophages in MaFIA animals alters animal activity and, in some studies, withdrawal thresholds^39,95^. In our neonatal behavior, we detected no changes in baseline withdrawal reflexes nor guarding scores due to the macrophage knockout (Fig.1C-D). However, we did observe that when we directly manipulated macrophage presence, there was an impact on withdrawal reflexes, but not spontaneous behaviors (Figs. 1D, 3E, and our previous work^5^). Since withdrawal thresholds are one of the most utilized behavioral readouts, it may be prudent to consider multiple behavioral assays when performing experiments depleting macrophages^5,21,39,95,96^. This confound is not a major concern for adolescent/adult behaviors, however, because the animal’s immune system had recovered before the second injury (Fig. 1). Similar concerns are also avoided when performing adoptive transfer experiments because we tested latent changes in primed macrophages, a major strength of this study.

We considered sex as a critical factor throughout our study. In our RNA-seq, we noticed some indications of differences between the sexes, especially in the early life incised condition. We do not have the power to critically evaluate these against one another, however, there are multiple lines of evidence that suggest the immune-modulated injury response differs between males and females^e.g.97^. In our evaluations in aged animals, with or without a BMT, we observed female-specific impact of the early life injury (Fig. 7). Transfer of HSCs alone was not sufficient to replicate the precise timing of hypersensitivity as if the host had been injured as a neonate, but primed HSC cells did produce a leftward shift of mechanical sensitivity, again only in females. Recently dos Santos et al found a delayed onset hypersensitivity in aged animals following an incision injury as well as blunted hyperalgesic priming in aged animals. This was connected with changes in inflammation in older rodents compared to younger animals^98^. We found a similar delay following a BMT in female incision at P147, but not in non-BMT males or females. Nevertheless, results reveal that the relatively minor neonatal hind paw incision injury causes “priming” of the bone marrow and resulted in a female specific long-lasting phenotype. Indeed, when evaluating the levels of p75NTR in our BMT animals, we found that isolated female peritoneal cells had significantly more p75NTR than males. These data, along with substantial literature on the heightened prevalence of chronic pain in females^99^, warrant additional studies that examine sex differences in injury-induced changes in immune cells over time to help better understand these phenomena.

Clinical studies have shown that neonatal injuries have the potential to cause a “primed” somatosensory system, and lead to long-term alterations in nociceptive processing^11,12,100,101^. The potential therapeutic ability of p75NTR inhibitors has been recently reviewed by Malik et al indicating a range of functional neurological disorders that are affected by p75NTR, while Norman and McDermott review potential therapeutic strategies in blocking p75NTR signaling for chronic pain^102,103^. New technologies that allow for the specific targeting of macrophage p75NTR could provide the specificity to prevent the priming effect on this cell type, while avoiding off-target effects of the nervous system^104^. Here, we discovered a change in the epigenetic landscape of macrophages due to early life incision that modulated neonatal nociceptive priming. Future studies designed to analyze the immune “pain memory” may lead to important therapeutic interventions for pediatric post-surgical pain.

## Methods

### Animals

Animals of both sexes were used for all studies. Neonatal animals were considered at postnatal day 7 (P7). Adolescent animals begun experimentation at P35-P42. Adult animals were analyzed at P147. Sex was analyzed as a critical biological factor for each experiment, and differences are noted in figure legends. In an effort to reduce stress, neonatal animals were kept with the dam when not actively analyzed behaviorally, and experiments were kept as short as possible (∼1 hour, never more than 1.5 hours). In all behavioral experiments, mice were acclimated to the room and were performed by the same experimenter under the same conditions including time of day, location, lighting and temperature. Animals were used +/-1 day from the indicated age. All animals were kept in an environment-controlled facility at Cincinnati Children’s Hospital Medical Center with free access to food and water while on a 14/10-hour light/dark cycle.

C57/Bl6 (CD45.2^+^) mice were bred in-house or purchased from Jax (Stock#: 000664). B6.SJL/BoyJ (CD45.1^+^) mice were bred in the CCHMC transplant core. Macrophage fas-induced apoptosis (MaFIA) animals were purchased from Jax (Stock#: 005070) and bred in-house. Myeloid/Macrophage reporter animals were generated by crossing the LysM-Cre positive animal (see Jax Stock#: 004781) with the tdTomato (tdTom) reporter mouse (B6.Cg-Gt(ROSA)26Sor^tm14(CAG-tdTomato)Hze^/J) purchased from The Jackson Laboratory (Stock#: 007914). Myeloid/Macrophage specific p75NTR knockout animals were generated by crossing the tamoxifen inducible LysM-CreERT2 animal (Jax 031674) to the p75NTRfl/fl animal that was kindly donated to us by Dr. Sung Yoon at the Ohio State University. All procedures were approved by the CCHMC Institutional Animal Care and Use Committee in compliance with AALAC approved practices.

### Behavioral Measures

Neonatal animals were transferred from their home cage to opaque chambers fitted with a translucent lid. They were acclimated for 10 minutes prior to any assessments. Adolescent animals were transferred from their home cage to raised chambers fitted with black side walls, translucent viewing front, back and lids, and a grid mesh bottom. These animals were acclimated for 30 minutes prior to any assessments. Behavioral data were gathered at neonatal baseline and one day (1d) following early life injury. Further data was gathered at adolescent baseline and 1d, 3d, 7d, 14d and 21d following adolescent injury. Behavioral assessments were performed as described before^5^ and below, and included a measure of spontaneous pain-like behavior as well as evoked muscle squeezing pain-like behavior.

Spontaneous pain-like behavior was measured as described previously by qualitatively assessing paw weight preference on a scale of zero to two, where zero is no guarding after injury, 1 indicates a shift in weight off of the injured paw, and 2 indicates a complete removal of the injured paw from the platform^5^. Assessments were made for a duration of 1 minute every 5 minutes for 30 minutes (neonates) or 1 hour (adolescents).

Muscle-directed withdrawal scores were measured using a digital pressure device (IITC Life Science Inc. Woodland Hills, CA, USA) with a dulled probe attachment ∼2 mm wide at the tip. In this assay, the animal was lightly scruffed and was held upright. One at a time, the hind paws were inserted between the two arms of the device. The top arm has a flat platform to hold the dorsal hindpaw in place, while the bottom has the dulled probe. Slowly, the medial plantar paw was pressed by the probe while the animal and force were observed. When the animal gave a robust withdrawal response, the force in grams was recorded. This was repeated for a total of three trials for each animal, separated by at least 5 minutes between trials for recovery. The maximum squeezing force implemented to a neonate was 150 grams and for an adolescent was 350 grams.

### Injections

To induce cell apoptosis in MaFIA animals, the designed drug AP20187 (AP; B/B Homodimerizer purchased from Takara #635058) was first prepared according to manufacturer’s directions in 100% ethanol. Prior to each experiment, fresh vehicle was prepared according to manufacturer’s directions for 10 mg/kg which contained 10% PEG-400, 86% Tween and 4% drug/vehicle. All animals were injected with a final volume of 30 µL by intraperitoneal (IP) injections. The drug or vehicle was added to the mixture immediately before injecting the animal. AP20187 has no known impact on wildtype (WT) animals and previous reports demonstrate that neonates injected at P1, P2, P3 and P7 show robust knockout of Csf1r expressing cells in tissue^41,105^. Tamoxifen (1000 μg at 10 mg/mL) was injected in all animals within a litter of potential LysM-CreERT2;p75NTRfl/fl genotype. The injection protocol began at P33 and was completed i.p. daily for 5 days. Animals injected with Evan’s Blue Dye (EBD, Sigma E2129-10) were injected as adolescence 16-20 hours prior to dissection with 200 µL of 1% EBD in sterile 0.9% saline.

### Surgical Hind Paw Incisions

Animals were anesthetized with 2-3% isoflurane. Once sufficiently anesthetized, animals were placed on their backs and their right hind paw was taped to a sterile cloth. The dorsal skin was disinfected using Chlorhaxiderm Scrub and 70% isopropyl alcohol. A longitudinal incision was made through the hairy skin lateral to the main saphenous innervation territory. The incision was continued in between the bones through to the body of the flexor digitorum brevis muscles. The muscles were then manipulated using blunt dissection techniques with #5 (neonate) and #3 (adolescent) forceps. The plantar skin was left untouched. The skin was sutured using 7-0 or 6-0 sutures for neonates and adolescents, respectively. Sham surgeries were performed exactly the same, except animals received only a suture through intact hairy hindpaw skin with no incision. Dual incision regimes were performed at both P7 and P35-P42, or P147 to the same hindpaw muscle.

### Dissections

For peripheral sites, the animal was heavily anesthetized with ketamine and xylazine (100 and 16 mg/kg, respectively) and the tissue was removed. This included hind paw muscle, tibia, potineal lymph node and spleen. For neural tissue, the animal was first cardiac perfused with ice-cold saline prior to dissection where ipsilateral DRGs were removed.

### Realtime PCR and Protein Arrays

RNA was isolated from BMDM cultures at the indicated time points using RNeasy Mini Kits (Qiagen Stock#: 74104). All isolation were performed exactly according to the manufacturer’s instructions. 500ng of RNA was reverse transcribed to cDNA and realtime PCR was performed using SYBR Green Master Mix on a StepOne realtime PCR system (Applied Biosystems). Quantitative PCR was analyzed by the ΔΔcycle threshold (CT) method with normalization to GAPDH. Differences in expression and standard error are determined from the normalized ΔΔCt. This is used to calculate fold change between conditions and values are then converted to a percent change where 2-fold = 100% change^106^. Cytokines from BMDMs were analyzed using proteome profilers (R&D Systems ARY006) loaded with 700 µLs of media. One membrane was removed due to a lack of positive control expression (siCON stim LPS+IFNy). Data was first normalized to the average positive control before percent change from experimental control was calculated.

### ATAC-Seq and RNA-seq

Peritoneal macrophages were isolated as indicated below from P7 naïve, P35 naïve, and P7 incised isolated at P35 animals. Samples were pooled across two animals of the same age and sex. Samples for ATAC-and RNA-Seq were processed by the sequencing core at CCHMC. Data were analyzed by the Bioinformatics Collaborative Services at CCHMC. Briefly, shared peaks between samples were obtained by merging peaks at 50% overlap with BEDtools v2.27.0 between replicates (peaks in all biological replicates) and treatment (merge at 50% all replicate peaks). These were then converted to Gene Transfer Format and peaks were counted with feature counts v1.6.2 (Rsubread package). Raw counts were normalized as transcripts per million and peaks were annotated by the nearest or overlapping gene and genomic features using an in-house script defined by: Promoter = 1kb upstream and downstream from peak; upstream = between 21 and 1kb from peak; priority = promoter>upstream>exon>intron. The R package DESeq2 v1.26.0 was used for differential analysis, with significance set to log2 fold change of 0.58 and FDR less than or equal to 0.05. RNA-seq was prepared similarly, where FASTQ formats were aligned to mm9 by STAR v2.6.le and stripped of duplicates by Sambamba v0.6.8. Read counts and differential expression were defined as before. Plots were generated by ggplot2 or PlotsReasy from CCHMC.

### Immunohistochemistry

Freshly dissected tissue was frozen in OCT medium on dry ice or snap frozen in liquid N_2_ for skeletal muscle and was then stored at-80°C. On a cryostat, tissue was sectioned at 20 µm and was melted onto a slide. Tissue was fixed in 4% paraformaldehyde, washed and blocked prior to primary antibody (WGA 1:500 (Invitrogen W32466); Dystrophin 1:250 (Abcam ab15277); p75NTR 1:1000 (Cell Signaling 82385); F4/80 1:500 (Abcam ab6640)) incubation overnight. The next day, the tissue was washed and stained with secondary antibodies before cover slipping and stained with DAPI (Fisher Scientific 17985-50.) For tissue that was genetically marked with a fluorescent dye, the tissue was post-fixed, washed, and mounted with media containing DAPI. Analysis was completed on Nikon’s NIS Elements where two-three nonconsecutive sections across two-three slides were averaged to create a score for each animal. Quantification of GFP+ macrophages in Fig. 1 were performed using Image J following similar procedures.

### Macrophage Isolation

*Peritoneal:* Animals were rapidly killed by cervical dislocation and cleaned with 70% ethanol. The peritoneum was injected with 1 mL (neonate) or 5 mLs (adolescent) of 3% fetal bovine serum (FBS) until bloated. The midsection was gently massaged to dislodge any cells, and then peritoneal fluid was extracted. Extracted peritoneal serum was processed for RNA extraction immediately, or sorted by the Flow Cytometry core at CCHMC by the LysMtdTom or MaFIA (GFP+) expression depending on the experiment and animal used. Cells were sorted in 2% FBS and into 50% FBS or PBS. Dead cells were marked with 7AAD and were removed from selection. Gating strategies were placed by the flow core and examples are presented.

*Bone marrow derived macrophages (BMDM):* Animals were prepared as above. The right leg was then detached from the body and placed into a dish containing HBSS. The femur and tibia were isolated, and the bone marrow was visualized by making horizontal cuts on either side of the knee. The bone marrow was flushed out of the bone with HBSS and a 22-gauge needle. HBSS with cells was collected, spun at 4°C and 1500 RPM for 5 minutes. The supernatant was discarded, and RPMI buffer (Catalog #11875093) supplemented with 10% fetal bovine serum (35-010-CV Lot 35010171) and gentamycin was added to resuspend the cells. GM-CSF (60 ng/mL) was added, and cells were incubated for 5 days at 37°C and 5% CO_2_. Additional media and GM-CSF was added halfway through the incubation. After maturation, cells were treated with siRNAs (Origene 10 ng/uL) and stimulants (NGF, BDNF 50 ng/uL or LPS+IFNγ 10 ng/uL each) as indicated in the figure legends. Knockdown efficiency tested across three duplexes, achieved 90% by PCR. Cells were stimulated for 24 hours before washing, trypsinizing, and/or scraping of the plates. Media was saved for protein array analyses. Cells were counted using a hemocytometer and then resuspended in buffer for RNA isolation using the RNeasy Mini Kit.

### Adoptive Transfer

Peritoneal macrophages from MaFIA animals were isolated as described above. ∼80k cells were sorted into PBS. These were pelleted and reconstituted in 10 μLs of sterile PBS. Naïve hosts were anesthetized and the right hindpaw was injected with all of the cells directly into either intact muscle, or the wound of a hindpaw incision before suturing. Animals were monitored following the transplant for adverse effects and behavioral responses.

### Bone Marrow Transplant

Tibia and femur bone marrow was collected using sterile tools from adolescent animals. Under a sterile hood, bones were flushed with PBS and treated with red blood cell lysis buffer until a transparent pellet was formed after centrifugation at 1100 RMP for 4 minutes at 4°C. After reconstitution, cells were passed through a 70 µm filter and were counted on a hemocytometer. One million cells were transferred to a naïve age and sex matched host that was previously exposed to radiation (administered in split doses, the first at 700 rads/700 cGy followed 3 hours later with a second dose of 4.75 Gy (475 rads/475 cGy)) to prepare for the transplant. Animals were monitored following the transplant in their immune compromised state. We successfully completed 28 out of 32 BMTs. The animals were allowed to recover for 16 weeks prior to further experimentation, to ensure a functional immune response after transplant^50,51^.

### Human induced pluripotent stem cell preparations and calcium imaging

Human induced pluripotent stem cells (iPSC) derived CD14+ monocytes (ATCC, Manassas, VA, Cat No. DYS0100) were plated in macrophage differentiation medium (RPMI-1640 Medium /10% Fetal Bovine Serum (FBS), supplemented with 100 ng/ml human M-CSF, and 1% Pen/Strep) in 24 well plate at a density of 10^5 cells/ml according to manufacturer’s directions. Monocytes were differentiated into macrophages within seven days. Half medium change and siRNA treatment were done at day three and day five, respectively. Briefly, 10 nM control (siCON), or siRNAs against p75NTR (sip75; Origene SR321107) were incubated with macrophages at day five in macrophage differentiation medium. At day six, macrophages treated with siRNAs were stimulated with either 50 ng/ml NGF or vehicle in macrophage differentiation medium for 24 hours. At day seven, the media was extracted and used for stimulation of the sensory neurons. Human iPSC-Derived Sensory Neuron Progenitors (AXOL; ax0555) were cultured on SureBond-XF-coated 96 well plate with Neural Plating Medium (AXOL, product codes: ax0053 and ax0033) for 24 hours as recommended by the manufacturer. Briefly, full media change was achieved by replacing the neural plating medium with sensory neuron maintenance medium supplemented with growth factors (25 ng/mL Glial-Derived Neurotrophic Factor (ax139855), 25 ng/mL Nerve Growth Factor (ax139789), 10 ng/mL Brain-Derived Neurotrophic Factor (ax139800), and 10 ng/mL Neurotrophin-3 (ax139811) and sensory maturation maximizer (ax0058) at a seeding density of 50000/cm^2^. Cells were treated with 2.5 µg/ml mitomycin C at day three after which the media was replaced with sensory neuron maintenance medium supplemented with growth factors and sensory maturation maximizer. Half media change was done at every three days for 21 days. All iPSC cultures and incubations were carried out in 5% CO_2_/ 37°C.

For imaging, cells were pretreated with 5 µg/ml Rhodamine (in DMSO, ThermoFisher R1244) for 30-45 minutes and further dosed with equal volume of siRNA treated and stimulated macrophage media. Neuronal activities were visualized using wide-field Nikon Ti2 inverted SpectraX with 10X LWD Phase high quality objective. To analyze changes in fluorescence, regions of interest (ROI) were randomly drawn around 15 cells across triplicate conditions. Calcium transients were measured for after stimulation with ATP (200μM) followed by capsaicin (1μM). KCl (300μM) was used as a positive control for cell viability. ΔF/FO were calculated where ΔF/FO = (Fmax – FO) / FO where Fmax: maximum intensity during stimulation and FO: the average intensity of immediately prior to stimulation.

### *Ex vivo* Preparation

This preparation has recently been described in detail^95^. In this study, pups were treated with either AP or vehicle as described above and were subjected to a P7 incision followed by a P35-P42 incision. Briefly, 7d after the second incision we dissected the animal HP muscle/tibial nerve/DRG/SC in continuity in ice cold oxygenated (95%O2/5% CO2) artificial cerebral spinal fluid (aCSF; 127.0 mM NaCl, 1.9mM KC l, 1.2 mM KH2PO4, 1.3 mM MgSO4, 2.4 mM CaCl2, 26.0 mM NaHCO3, and 10.0 mM D-glucose: O2aCSF). The preparation was transferred to a new dish where the muscle and DRGs were isolated in separate circulating baths of O2aCSF. The temperature was slowly raised to 32°C. In L3 or L4 an electrode was filled with 5% neurobiotin and 1M potassium acetate for sharp electrode single unit recordings using Quartz electrodes (impedance>150MΩ). A suction electrode was placed on the side of the tibial nerve to send search stimuli. An impaled cell was identified by the presence of an action potential (AP) recording from the search stimulus. A receptive field on the muscle was then located using a concentric bipolar electrode. If found, the cell was counted in our analyses and mechanical, thermal and chemical stimuli were applied in that order. First von Frey fiber thresholds were identified using fibers from 0.07-10g of force applied for ∼1-2 seconds. If the cell did not respond to the maximum, physical push was applied to determine if the cell would respond to any mechanical force. Next ice-cold saline (∼2°C) was applied to the circulating bath over the receptive field, followed by hot ∼53°C saline. After washout, a low concentration mixture of lactic acid, protons and ATP (applied just before stimulation) were oxygenated and flooded the muscle chamber through an in-line heater for 2 minutes (15 mM lactate, 1μM ATP, pH 7.0). This was repeated with a high concentration of the same metabolites (50 mM lactate, 5 μM ATP, pH 6.6). Mechanical and thermal responses were then repeated.

Activity was recorded by the Spike2 program (Cambridge Electronic Design) and was analyzed offline. At the end of the preparation the nerve length from suction electrode to DRG was measured to calculate conduction velocity (CV), where a Group IV afferent was defined as being (≤1.2 m/s) or Group III afferents (1.2–14 m/s). Mechanical thresholds were set at the minimum force required to evoke an action potential, firing rate (FR) was calculated as the maximum number events that occurred over a 200 ms bin, and instantaneous frequency (IF) was calculated to determine the maximum response to a stimulus. Distribution analyses were completed to determine if there was a shift in the number of responders in our conditions.

### Statistical Analyses

Data was analyzed using SigmaPlot software (v14.5). Critical significance value was set to α<0.05. All data were first checked for normality by Shapiro-Wilk to determine parametric or nonparametric tests. Individual tests used are described in the figure legends. Behavioral data of the same animal over time and between groups were analyzed using a two-way repeated measures (RM) ANOVA. In situations in which different animals were compared across multiple time points and groups, such as PCR data, a two-way ANOVA was used. Data at a single time points across multiple groups including much of the electrophysiological data were analyzed by a one-way ANOVA. Proportions data were analyzed using a Chi-squared or Fisher’s exact test as marked in the figure legends. Graphs were made using GraphPad Prism (v9). In all experiments in which there is a potential for bias including behavior, electrophysiology, cell culture and IHC the investigator was blinded to the conditions. For behavior, animals were assigned to groups by a random number generator. Biological and technical replicates were completed for each experiment and are appropriately marked in the figure legends.

## Author contributions

A.J.D. and M.P.J. conceived and designed the study and wrote the manuscript. A.J.D. completed most of the experiments and collected most of the data. A.J.D. performed the analyses with input from M.P.J. A.F. completed the experiments at P147 and the human culture experiments. A.M.W. completed various experiments including some IHC and qPCR. A.P. completed bioinformatics data processing. L.F.Q. assisted in ex vivo recordings and interpretations of data. G. D. and H.E. facilitated experiments utilizing bone marrow isolation, differentiation and stimulation of macrophages. G.D. helped with the conceptualization of the study. D. L. and B.W. assisted with the BMT experiment planning and execution. O.D., C.F., M.T.W., and L.C.K. helped perform ATAC-seq and RNA-seq processing. M.C.H. managed the colony and performed genotyping assays.

## Acknowledgements

We would like to recognize others who have provided support for this project including: The Research Flow Cytometry Core in the Division of Rheumatology and the Confocal Imaging Core at CCHMC; Dr. Yoon at the Ohio State University who provided to us the p75NTRfl/fl animals; The Center for Autoimmune Genomics and Etiology (CAGE) at CCHMC and; The Comprehensive Mouse and Cancer Core for their assistance in the completion of the bone marrow transplants and care for the animals. This work was supported by grants from the NIH to M.P.J. (R01NS105715 and R01NS113965), A.J.D. (F31NS122494), D.L. (R01 HL160614), L.C. K. (P30 AR070549), and a CCHMC ARC award (#53632) to L.C.K. and M.T.W.. D.L. is a scholar of the Leukemia and Lymphoma Society.

