## Supplementary Figures for "Macrophage epigenetic memories of early life injury drive neonatal nociceptive priming"

#### Supplemental Figure 1

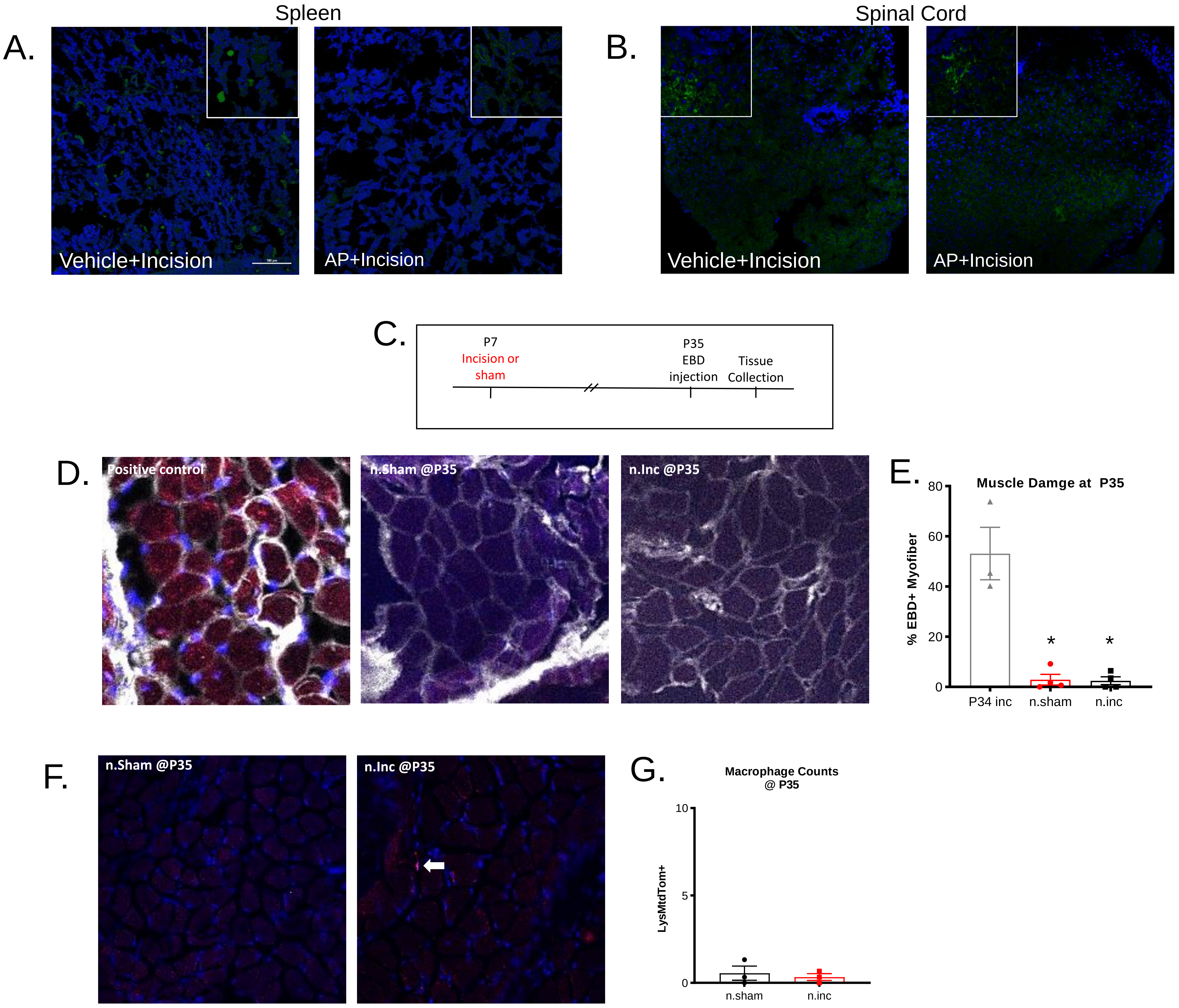

### Supplemental Figure 2

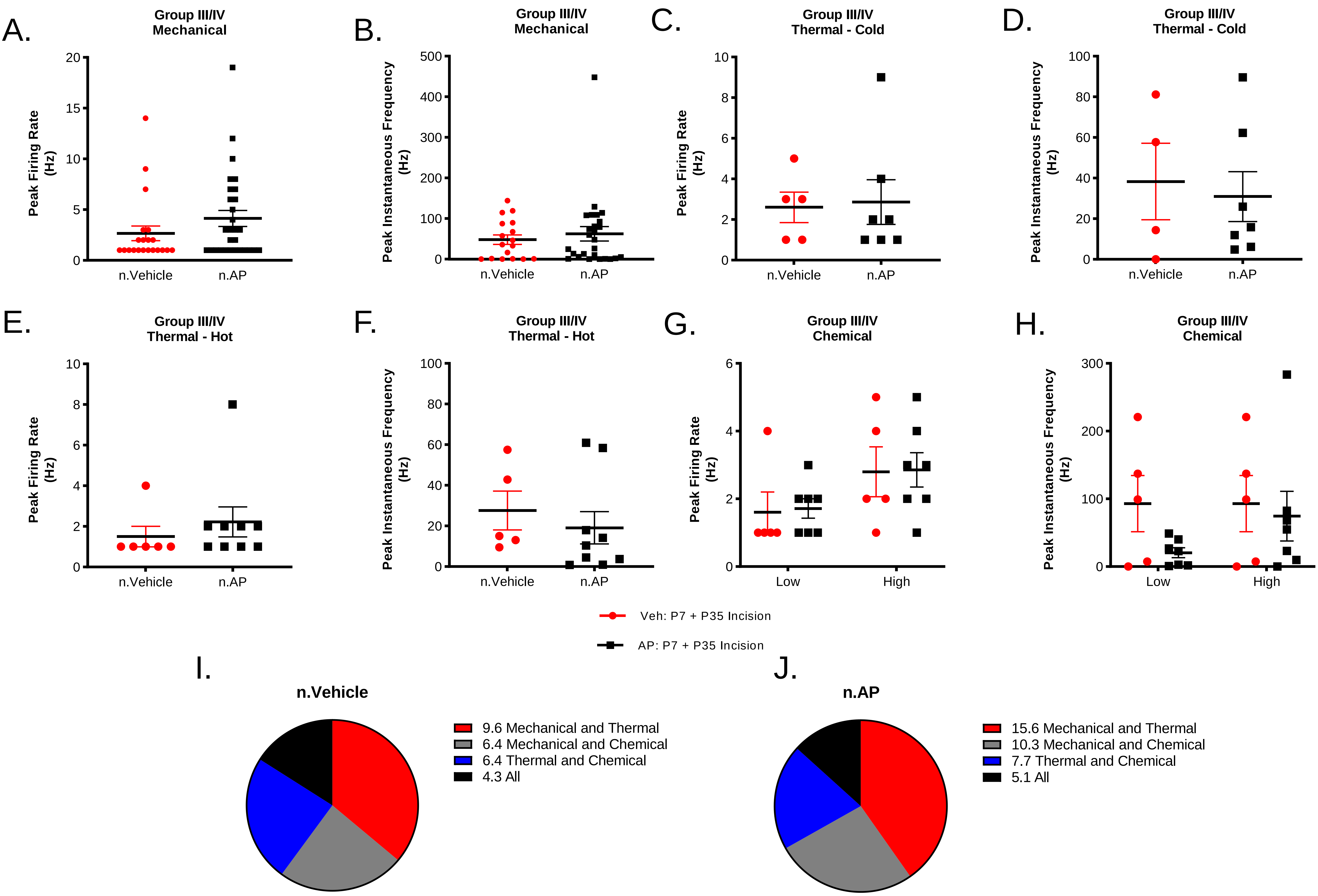

#### Supplemental Figure 3

A.

| Condition | ATAC-seq peak count | Most similar cell type (public data) | Most similar molecule (public data) |
|---|---|---|---|
| P7 Naïve Isolation | 76,624 | BMDM_macrophages | Pu1 |
| P7 Naïve Isolation | 80,621 | BMDM_macrophages | Pu1 |
| P7 Naïve Isolation | 58,325 | BMDM_macrophages | Pu1 |
| P35 Naïve Isolation | 75,182 | BMDM_macrophages | Pu1 |
| P35 Naïve Isolation | 76,673 | macrophage | Pu1 |
| P35 Naïve Isolation | 74,305 | macrophage | Pu1 |
| P7 Incised, P35 Isolation | 57,259 | macrophage | Pu1 |
| P7 Incised, P35 Isolation | 56,885 | Macrophage | Pu1 |
| P7 Incised, P35 Isolation | 69,108 | BMDM_macrophages | Pu1 |

B.

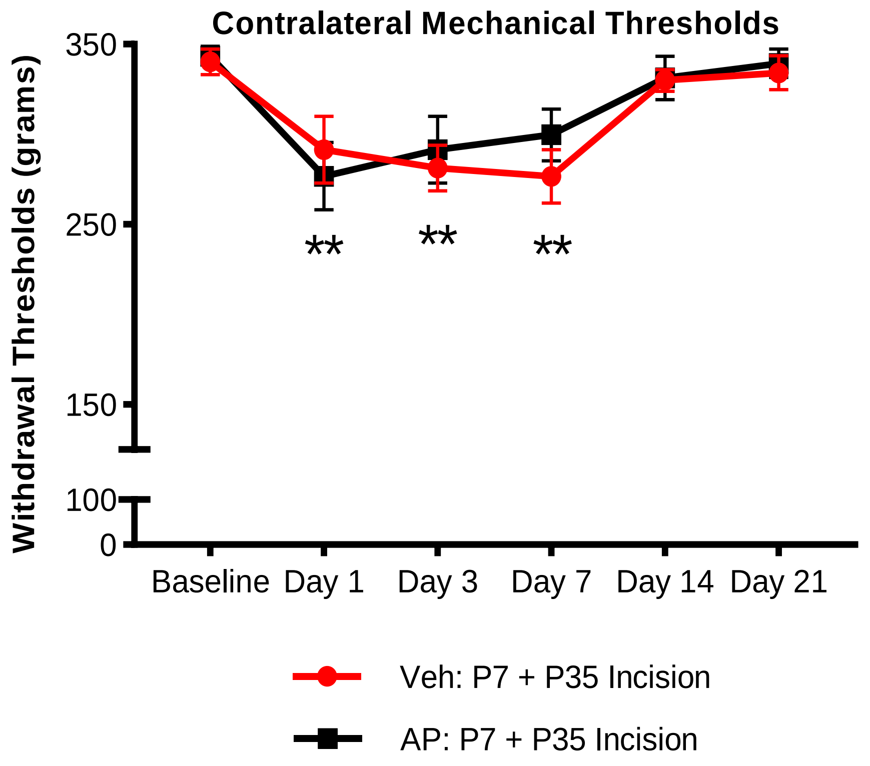


C.

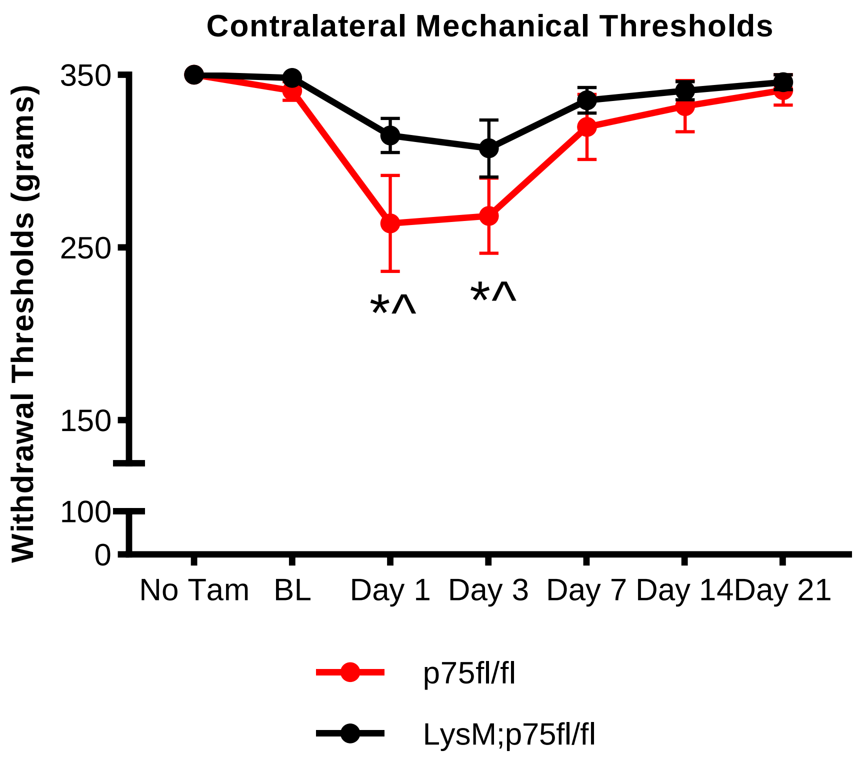

### Supplemental Figure 4

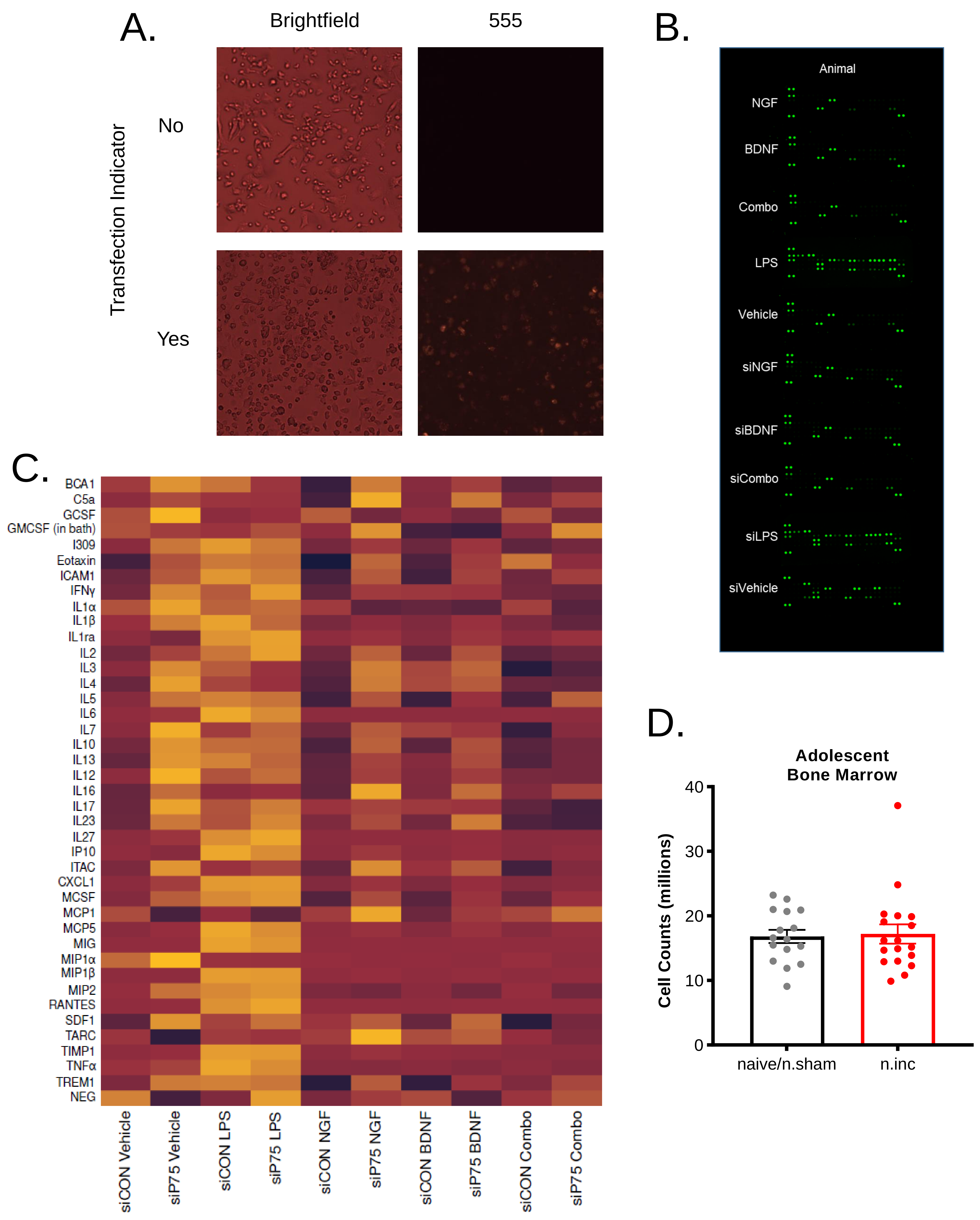

### Supplemental Figure 5

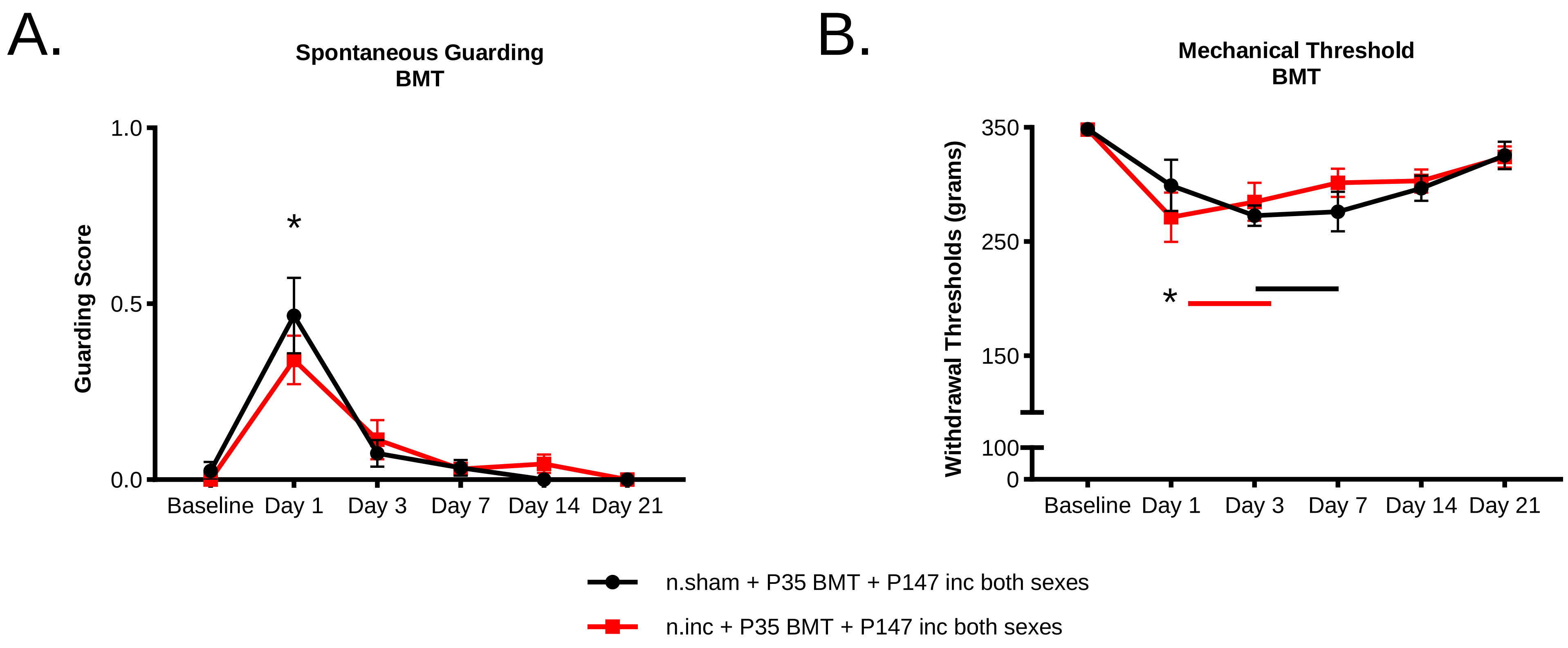

### Supplemental Table

| Protein | sip75+Veh | siCON+LPS | sip75+LPS | siCON+NGF | sip75+NGF | siCON+BDNF | sip75+BDNF | siCON+Combo | sip75+Combo |
|---|---|---|---|---|---|---|---|---|---|
| | % Change from siCON+Vehicle | | | | | | | | |
| BCA1 | 10 ± 17 % | 47 ± 13 % | 0 ± 10 % | -20 ± 4 % | 11 ± 14 % | -7 ± 9 % | -13 ± 6 % | -7 ± 11 % | -24 ± -24 % |
| C5a | 12 ± 15 % | 32 ± 21 % | 19 ± 17 % | -43 ± 4 % | 160 ± 72 % | -2 ± 11 % | 95 ± 45 % | 1 ± 20 % | 18 ± 18 % |
| GCSF | 1258 ± 733 % | 136 ± 112 % | 290 ± 164 % | 18 ± 11 % | 30 ± 54 % | 0 ± 27 % | 14 ± 40 % | 4 ± 3 % | 27 ± 27 % |
| GMCSF | -18 ± 26 % | 36 ± 51 % | 30 ± 42 % | 5 ± 26 % | 31 ± 33 % | -26 ± 11 % | -41 ± 7 % | 7 ± 29 % | 30 ± 30 % |
| I309 | 127 ± 66 % | 482 ± 284 % | 252 ± 116 % | 8 ± 24 % | 37 ± 42 % | -7 ± 23 % | 33 ± 37 % | -12 ± 20 % | -13 ± -13 % |
| Eotaxin | 57 ± 33 % | 219 ± 65 % | 189 ± 53 % | -18 ± 6 % | 105 ± 18 % | 14 ± 15 % | 59 ± 18 % | 217 ± 62 % | 27 ± 27 % |
| ICAM1 | 58 ± 49 % | 286 ± 162 % | 199 ± 66 % | -16 ± 15 % | 76 ± 48 % | -20 ± 41 % | 44 ± 43 % | 24 ± 32 % | 43 ± 43 % |
| IFNγ | 76 ± 10 % | 158 ± 31 % | 177 ± 39 % | -1 ± 14 % | 14 ± 16 % | 36 ± 13 % | 9 ± 14 % | 14 ± 9 % | -17 ± -17 % |
| IL1a | 575 ± 472 % | 665 ± 437 % | 427 ± 176 % | 4 ± 34 % | -15 ± 38 % | -15 ± 46 % | -16 ± 40 % | 2 ± 27 % | -25 ± -25 % |
| IL1b | 253 ± 153 % | 734 ± 575 % | 322 ± 154 % | 2 ± 21 % | 34 ± 39 % | 12 ± 29 % | 53 ± 48 % | 21 ± 27 % | -25 ± -25 % |
| IL1ra | -31 ± 36 % | 669 ± 167 % | 4104 ± 3619 % | 30 ± 16 % | 193 ± 189 % | 5 ± 30 % | 240 ± 216 % | 5 ± 12 % | 245 ± 245 % |
| IL2 | 29 ± 10 % | 243 ± 86 % | 295 ± 84 % | 7 ± 20 % | 102 ± 26 % | -4 ± 10 % | 84 ± 34 % | -9 ± 9 % | -15 ± -15 % |
| IL3 | 11 ± 16 % | 40 ± 7 % | 7 ± 7 % | 0 ± 12 % | 17 ± 3 % | 19 ± 14 % | 8 ± 14 % | -7 ± 11 % | -20 ± -20 % |
| IL4 | 140 ± 133 % | 63 ± 16 % | 35 ± 14 % | -6 ± 9 % | 98 ± 33 % | 76 ± 69 % | 60 ± 20 % | 17 ± 20 % | -12 ± -12 % |
| IL5 | 28 ± 15 % | 136 ± 75 % | 103 ± 60 % | -15 ± 14 % | 31 ± 31 % | -16 ± 18 % | 1 ± 20 % | -13 ± 11 % | 56 ± 56 % |
| IL6 | 1036 ± 920 % | 35132 ± 18092 % | 19944 ± 9191 % | -52 ± 14 % | 31 ± 44 % | -23 ± 20 % | 112 ± 128 % | 65 ± 69 % | 13 ± 13 % |
| IL7 | 31 ± 13 % | 28 ± 30 % | 37 ± 4 % | 3 ± 15 % | 13 ± 24 % | 28 ± 30 % | -7 ± 12 % | -7 ± 13 % | -8 ± -8 % |
| IL10 | 91 ± 30 % | 148 ± 75 % | 113 ± 38 % | -8 ± 15 % | 57 ± 33 % | -1 ± 21 % | 42 ± 23 % | 2 ± 16 % | -6 ± -6 % |
| IL13 | 62 ± 25 % | 128 ± 74 % | 86 ± 31 % | 14 ± 23 % | 22 ± 22 % | 24 ± 22 % | 30 ± 21 % | 9 ± 17 % | -3 ± -3 % |
| IL12 | 92 ± 36 % | 70 ± 50 % | 96 ± 36 % | -7 ± 23 % | 9 ± 19 % | 3 ± 19 % | 7 ± 16 % | 5 ± 13 % | -19 ± -19 % |
| IL16 | 15 ± 25 % | 49 ± 50 % | 21 ± 19 % | 23 ± 32 % | 63 ± 47 % | 23 ± 47 % | 40 ± 52 % | 36 ± 38 % | 16 ± 16 % |
| IL17 | 93 ± 56 % | 108 ± 79 % | 110 ± 29 % | 48 ± 34 % | 27 ± 27 % | 45 ± 35 % | 17 ± 29 % | 12 ± 14 % | -20 ± -20 % |
| IL23 | 35 ± 17 % | 64 ± 24 % | 116 ± 24 % | 21 ± 19 % | 20 ± 12 % | 6 ± 2 % | 69 ± 24 % | 3 ± 17 % | -24 ± -24 % |
| IL27 | 67 ± 22 % | 1327 ± 627 % | 1556 ± 336 % | 17 ± 15 % | 0 ± 7 % | 25 ± 16 % | 20 ± 3 % | -2 ± 6 % | -21 ± -21 % |
| IP10 | -45 ± 22 % | 1977 ± 118 % | 2556 ± 1661 % | 3 ± 17 % | 202 ± 132 % | 8 ± 15 % | 112 ± 124 % | -22 ± 6 % | 61 ± 61 % |
| ITAC | 34 ± 28 % | 23 ± 17 % | 29 ± 19 % | 4 ± 16 % | 43 ± 9 % | 16 ± 18 % | 17 ± 4 % | -4 ± 11 % | -10 ± -10 % |
| CXCL1 | 48 ± 22 % | 794 ± 388 % | 660 ± 232 % | -4 ± 17 % | 33 ± 24 % | -19 ± 12 % | 10 ± 17 % | -3 ± 18 % | -9 ± -9 % |
| MSCF | 35 ± 32 % | 207 ± 102 % | 183 ± 72 % | -13 ± 16 % | 32 ± 25 % | -12 ± 17 % | 9 ± 27 % | 2 ± 17 % | 10 ± 10 % |
| MCP1 | -35 ± 27 % | 31 ± 32 % | -16 ± 27 % | 9 ± 17 % | 37 ± 12 % | -20 ± 15 % | -13 ± 17 % | 18 ± 26 % | 25 ± 25 % |
| MCP5 | 36 ± 29 % | 2115 ± 925 % | 1624 ± 481 % | 41 ± 4 % | 75 ± 9 % | -13 ± 11 % | 82 ± 17 % | -3 ± 13 % | 39 ± 39 % |
| MIG | 37 ± 24 % | 13556 ± 6039 % | 10033 ± 2898 % | 13 ± 22 % | 21 ± 19 % | 8 ± 22 % | 10 ± 13 % | 6 ± 13 % | -18 ± -18 % |
| MIP1a | 1670 ± 1601 % | 165 ± 144 % | 85 ± 73 % | -9 ± 31 % | 8 ± 37 % | -18 ± 27 % | -14 ± 29 % | -6 ± 29 % | -20 ± -20 % |
| MIP1b | -9 ± 31 % | 7235 ± 2620 % | 8906 ± 4543 % | 79 ± 35 % | 129 ± 55 % | 34 ± 25 % | 105 ± 71 % | 5 ± 5 % | 88 ± 88 % |
| MIP2 | 4924 ± 3763 % | 14064 ± 7228 % | 11931 ± 4206 % | 18 ± 37 % | 10 ± 37 % | 24 ± 23 % | -6 ± 32 % | 42 ± 19 % | -19 ± -19 % |
| RANTES | 59 ± 31 % | 11986 ± 3798 % | 12424 ± 1735 % | 19 ± 16 % | 1 ± 19 % | -24 ± 13 % | 9 ± 19 % | 31 ± 10 % | 3 ± 3 % |
| SDF1 | 42 ± 20 % | 50 ± 5 % | 66 ± 25 % | 34 ± 13 % | 25 ± 5 % | 8 ± 9 % | 38 ± 7 % | -2 ± 4 % | -5 ± -5 % |
| TARC | -56 ± 25 % | 4194 ± 4177 % | 4434 ± 4455 % | 3034 ± 3026 % | 6719 ± 6648 % | 4422 ± 4411 % | 4978 ± 4979 % | 1351 ± 1351 % | 1934 ± 1934 % |
| TIMP1 | -6 ± 10 % | 1159 ± 345 % | 1097 ± 305 % | 7 ± 15 % | 42 ± 11 % | 16 ± 8 % | -9 ± 12 % | -3 ± 15 % | -25 ± -25 % |
| TNFa | 572 ± 469 % | 13303 ± 7767 % | 8392 ± 5043 % | 0 ± 50 % | 24 ± 54 % | -28 ± 21 % | -30 ± 29 % | 2 ± 32 % | -26 ± -26 % |
| TREM1 | 34 ± 30 % | 137 ± 90 % | 88 ± 39 % | -12 ± 19 % | 33 ± 32 % | -11 ± 23 % | 10 ± 36 % | 21 ± 17 % | 34 ± 34 % |
