## Supplementary Figure Legends for "Macrophage epigenetic memories of early life injury drive neonatal nociceptive priming"

**Supplemental Figure 1. No persistent effects of early life incision are found on muscle integrity.**

**A.** Representative images from the spleen at P8 after treatment with either vehicle or AP in MaFIA animals indicating a loss of GFP+ cells in the AP treated group. **B.** Images of the spinal cord of the same animals indicating continued expression of GFP in both groups. **C.** Experimental design for the collection of tissue in D-G. **D.** Representative images of EBD (red) counter stained with WGA (white) and DAPI (blue) in acutely injured tissue compared to latent changes four weeks after an early life incision or sham. **E.** In tissue damaged two days before dissection, there is robust EBD leakage into myofibers that is absent in neonatal injured myofibers (*p<0.001 vs. P34 inc., One-way ANOVA, Tukey’s, n=3-4/group). **F.** Images indicating LysM+ myeloid cells (tdTomato reporter) in adolescent tissue from neonatal injured or sham animals. **G.** There is no difference in the number of macrophages present in the muscle in neonatal incised animals compared to sham operated animals (p=0.643 n.inc vs. n.sham, One-way ANOVA, Tukey’s, n=3/group). Data shown as mean +/- SEM.

**Supplemental Figure 2. Additional electrophysiological properties of DRG neurons after repeated surgical injury.**

**A-B.** There is no change in mechanical firing rate (A, p=0.135) nor instantaneous frequency (B, p=0.804) between groups. **C-D.** Firing rates and instantaneous frequencies of cold responders is not different between groups (FR: p=0.876, IF: p=0.738, n.AP vs. n.Vehicle, n=7 and 5 respectively, ANOVA on Ranks, Dunn’s). **E-F.** There are no differences in heat responders between groups (FR: p=0.226, IF: p=0.386, n.AP vs. n.Vehicle, n=9 and 6 respectively, ANOVA on Ranks, Dunn’s). **G-H.** There is no difference in the firing rate nor instantaneous frequencies of chemically sensitive cells (Low: FR: p=0.530, IF: p=0.343, n.AP vs. n.Vehicle, n=7 and 5 respectively, ANOVA on Ranks, Dunn’s, High: FR: p=0.948, IF: 0.748, n.AP vs. n.Vehicle, n=7 and 5 respectively, ANOVA on Ranks, Dunn’s). **I-J.** There is no change in the proportion of polymodal responders between groups (n.AP vs. n.Vehicle, Chi-square or Fisher’s Exact). Data shown as mean +/- SEM.

**Supplemental Figure 3. ATAC-seq quality control and the effect of dual incision on contralateral mechanical withdrawal thresholds.**

**A.** Quality control analyses reveal a pure population of cells that are likely macrophages based on open chromatin regions and previous sequencing datasets. **B.** Animals that received dual incision regardless of treatment with AP or vehicle had a significant reduction of contralateral mechanical withdrawal thresholds (F=15.337) that lasted for at least seven days (**p<0.05 vs. BL, both groups, Two-way RM ANOVA, Tukey’s, n=10-12. **C.** Contralateral muscle squeezing is affected by the knockout of p75NTR in LysM+ cells following dual incision (F=13.047) on days one and three (*p<0.001 vs. p75fl/fl BL, ^p<0.05 vs. LysM;p75fl/fl, Two-way RM ANOVA, Tukey’s, n=7/group). Data shown as mean +/- SEM.

**Supplemental Figure 4: Transfection efficiency and regulation of bone marrow derived macrophages.**

**A.** Example pictures of the transfection efficiency to knockdown p75NTR in BMDMs. **B.** Example blots macrophages that were treated with different stimulants and the knockdown of p75NTR. **C.** A heatmap consisting of all manipulations for the 40 proteins included in the arrays. **D.** An analysis indicating no difference in the number of cells obtained from the bone marrow of early life incised or naïve or sham animals isolated at P35 (p=0.617 controls vs. n.inc, ANOVA on Ranks, n=16-18/group). “LPS” indicates LPS+IFNγ stimulation, “si” indicates treatment with the siRNA against p75NTR, “Combo” indicates both NGF and BDNF stimulation applied. Data shown as mean +/- SEM.

**Supplemental Figure 5: Sex aggregated behavioral results following neonatal injury, BMT and re-injury.**

**A**. After recovery from the BMT, there is a main effect of day (F=25.051). Animals display no guarding behaviors prior to incision. Following an incision both n.sham and n.inc groups from both sexes guarded for one day only (*p<0.01 vs. BL, Two-way RM ANOVA, Tukey’s). **B.** There was also an effect of day on withdrawal thresholds (F=9.099) that was significant at days one and three in the primed group but was delayed in controls with significant reductions at days three and seven (*p<0.01 vs. BL, Two-way RM ANOVA, Tukey’s, n=10-11/group).

**Supplementary Table: Quantification of protein arrays.**
